# Epigenetic priming of embryonic lineages in the mammalian epiblast

**DOI:** 10.1101/2024.01.11.575188

**Authors:** Miquel Sendra, Katie McDole, Daniel Jimenez-Carretero, Juan de Dios Hourcade, Susana Temiño, Morena Raiola, Léo Guignard, Philipp J Keller, Fátima Sánchez-Cabo, Jorge N. Domínguez, Miguel Torres

## Abstract

Understanding the diversification of mammalian cell lineages is an essential to embryonic development, organ regeneration and tissue engineering. Shortly after implantation in the uterus, the pluripotent cells of the mammalian epiblast generate the three germ layers: ectoderm, mesoderm and endoderm^1^. Although clonal analyses suggest early specification of epiblast cells towards particular cell lineages^2–4^, single-cell transcriptomes do not identify lineage-specific markers in the epiblast^5–11^ and thus, the molecular regulation of such specification remains unknow. Here, we studied the epigenetic landscape of single epiblast cells, which revealed lineage priming towards endoderm, ectoderm or mesoderm. Unexpectedly, epiblast cells with mesodermal priming show a strong signature for the endothelial/endocardial fate, suggesting early specification of this lineage aside from other mesoderm. Through clonal analysis and live imaging, we show that endothelial precursors show early lineage divergence from the rest of mesodermal derivatives. In particular, cardiomyocytes and endocardial cells show limited lineage relationship, despite being temporally and spatially co-recruited during gastrulation. Furthermore, analysing the live tracks of single cells through unsupervised classification of cell migratory activity, we found early behavioral divergence of endothelial precursors shortly after the onset of mesoderm migration towards the cardiogenic area. These results provide a new model for the phenotypically silent specification of mammalian cell lineages in pluripotent cells of the epiblast and modify current knowledge on the sequence and timing of cardiovascular lineages diversification^1^^2,13^.

## Introduction

A fundamental question in developmental biology is how embryonic cell lineages are specified and differentiate towards specific fates required for the construction of tissues and organs. The epiblast of the early mammalian embryo contains pluripotent cells able to contribute to all cell lineages of the new organism. Shortly after implantation in the uterus, the pluripotent epiblast cells differentiate into the definitive germ layers: endoderm, mesoderm, and ectoderm. When gastrulation begins, posterior epiblast cells gradually delaminate, forming the so-called primitive streak, and differentiate into mesodermal cells that migrate towards the anterior pole. Mesodermal progenitors thus relocate during gastrulation to distinct embryonic regions, whereas the specific signals they sense during migration and at destination steer their differentiation into particular cell types and organs^1,14,15^.

Despite the pluripotent nature of epiblast cells^16^ clonal analyses suggest that they initiate specification shortly before gastrulation, segregating progenitors of the yolk sac endothelium versus blood^17^, myocardium versus endocardium^3,4^ and definitive endoderm versus anterior mesoderm^18^. These observations suggest the presence of molecular diversity within the epiblast at embryonic day (E) 6.5. Contrary to this notion, single-cell and spatial RNA sequencing (scRNAseq) of the E6.5 epiblast identifies signatures linked to gastrulation priming, but fails to identify any cell lineage-specific expression profile^5–11^.

In addition to RNA expression, epigenetic modifications that influence chromatin accessibility contribute to cellular diversity^6,19^ and predicts the developmental competence of cell progenitors^20^. Notably, in the E8.25 mouse embryo, distinct open chromatin region sets associate with transcriptionally defined cell progenitor populations, fated to different lineages, like the erythroid or cardiac lineages^5,21^. Given the functional importance of chromatin remodelling complexes in gastrulation^22^, we posited that chromatin accessibility analysis could offer insights into inferring developmental trajectories in early organogenesis. Here, we used single-cell epigenomics to identify sub-types of pre-gastrulation epiblast cell populations by their epigenetic profile. Three clusters stood out by showing clear signatures of mesodermal/endothelial, endodermal and neuroectodermal fates, respectively. To understand the functional relevance of these findings, we applied clonal analysis and live imaging to further investigate the specification and differentiation of the embryonic endothelial cell lineage. We report that endothelial precursors transit through a specified but differentiation-silent phase that extends through gastrulation and during which they are indistinguishable in behaviour and spatio- temporal distribution from other mesodermal cells, including cardiomyocyte precursors. Shortly after their ingression through the primitive streak and during their migration towards their definitive position in the cardiogenic region, endothelial precursors show the first signs of differentiation from the rest of mesodermal cells by exhibiting specific migratory activity and increased affinity for the endoderm. These results show epigenetic priming of the endothelial fate in the epiblast, followed by early differentiation of endothelial/endocardial precursors shortly after gastrulation.

## Results

### Epigenetic priming of embryonic lineages in the mouse epiblast

To discern whether the E6.5 epiblast contains cells primed for specific fates, we reanalysed scRNAseq data from the mouse gastrulation atlas^5^. For this, we studied the emergence of expression signatures of the differentiated E8.5 cell types at earlier time points using UCell^23^. We observed no specific signatures rising above noise until E7.0, when an endothelial signature was detected (Endothelium, **Fig. S1_1-S1_4**). This may correspond to precursors of endothelial cells in the yolk sac and endocardium, in line with the detection of transcriptional differences between cardiomyocyte and endocardial progenitors at E7.25 by scRNAseq^24^ (**Fig. S1_5**). These data confirmed the absence of a transcriptomic signature for embryonic mesodermal lineages before gastrulation.

Next, we studied the epigenetic status of pre-gastrula cells by generating chromatin accessibility profiles from E6.5 mouse embryos using single-nucleus assay for transposase accessible chromatin (snATACseq). We isolated 38 embryos that lacked visual signs of primitive streak or nascent mesoderm formation (**Fig. 1A)**. After discarding the extraembryonic portion, we obtained 13,750 nuclei for snATAC-seq. Following sequencing and quality control, 7,283 nuclei were annotated by examining the gene start sites associated to the open-chromatin regions detected and correlating them to the marker genes of the cell types present at E6.5, as defined by^5^. This annotation identified 5778 epiblast, 1370 endoderm and 129 extraembryonic ectoderm nuclei (**Fig. 1B**).

**Figure 1:**
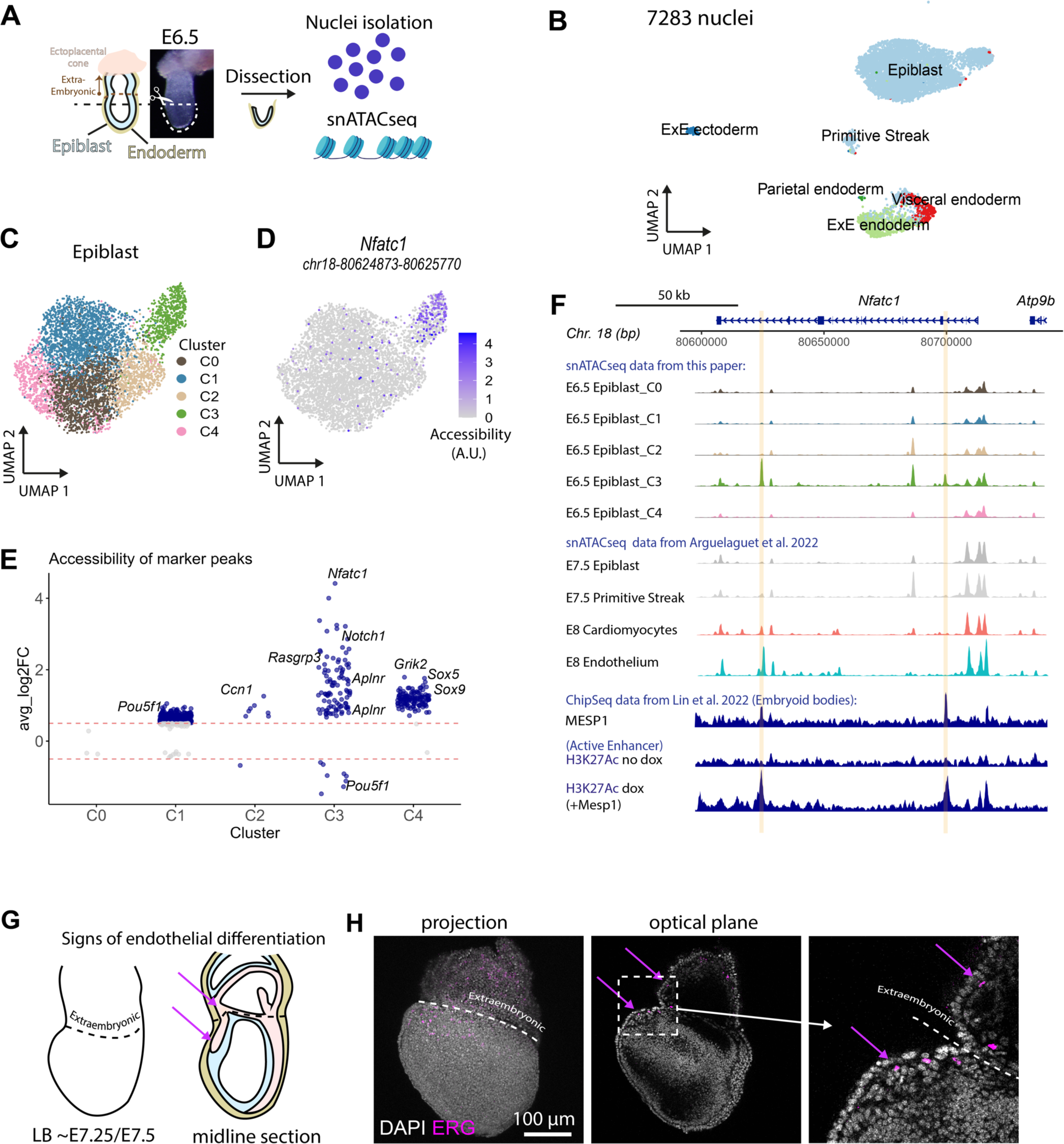
Single nucleus chromatin accessibility profiling reveals endocardial priming in the E6.5 epiblast. (A) Experimental setup. (B) UMAP representation of nuclei annotated with Mouse Gastrulation E6.5 reference^5^ (n = 7283 nuclei from 38 pooled embryos). (C) Epiblast subset classified by a shared nearest neighbour based clustering algorithm (n = 5573 nuclei from 38 pooled embryos). (D) Accessibility of Cluster 3 (C3) marker region belonging to an *Nfatc1* putative intronic enhancer (chr18:80624873-80625770). (E) Differentially accessible regions in epiblast clusters. A Log2Fold Change line is depicted at -0.5 and 0.5. (F) Coverage plot of the whole *Nfatc1* regulatory region^50^. The first five rows correspond to pseudobulk ATAC- seq signal for E6.5 epiblast clusters. The following four rows show pseudobulk ATAC-seq signal for revelant cell types at later time points^51^. Notice how E6.5 epiblast C3 shows the same accessibility pattern as the differentiated endothelium (E8). Last three rows represent ChipSeq tracks for Mesp1 and enhancer histone marks done in embryoid bodies^27^. (G) Schematics of H experimental setup. (H) ERG immunostaining revealing endo- thelial precursors both in the embryo proper and extraembryonic regions (n = 7 embryos).

We then specifically analysed epiblast nuclei, which identified five clusters (**Fig. 1C**). Notably, cluster 3 showed the strongest pattern of differentially open chromatin regions. Furthermore, several of the more differentially activated regions in Cluster 3 associated with endothelial markers, including the early endocardium marker *Nfatc1* ^25,26^; *Notch1*, which reports the earliest bias towards endocardium in cardiac progenitors at E7.25^24^ and other endothelial cell markers (**Fig. 1D-F**, supp. data 1). Integration of ChipSeq data from embryoid bodies^27^ revealed that 32% of cluster 3 and 12% of cluster 2 marker peaks acquired the H3K27Ac enhancer activation mark upon the induction of the master mesoderm regulator *Mesp1*, whereas other clusters did not show this association (**Fig. 2A**). Using chromVAR, we then assessed the accessibility of transcription factor DNA-binding motifs across epiblast clusters^28^. The top enriched motifs— FOX family in cluster 2, GATA in cluster 3, and SOX in cluster 4— relate to definitive endoderm, nascent mesoderm, and neural ectoderm lineages, respectively^7^ (**Fig. 2B**).

**Figure 2:**
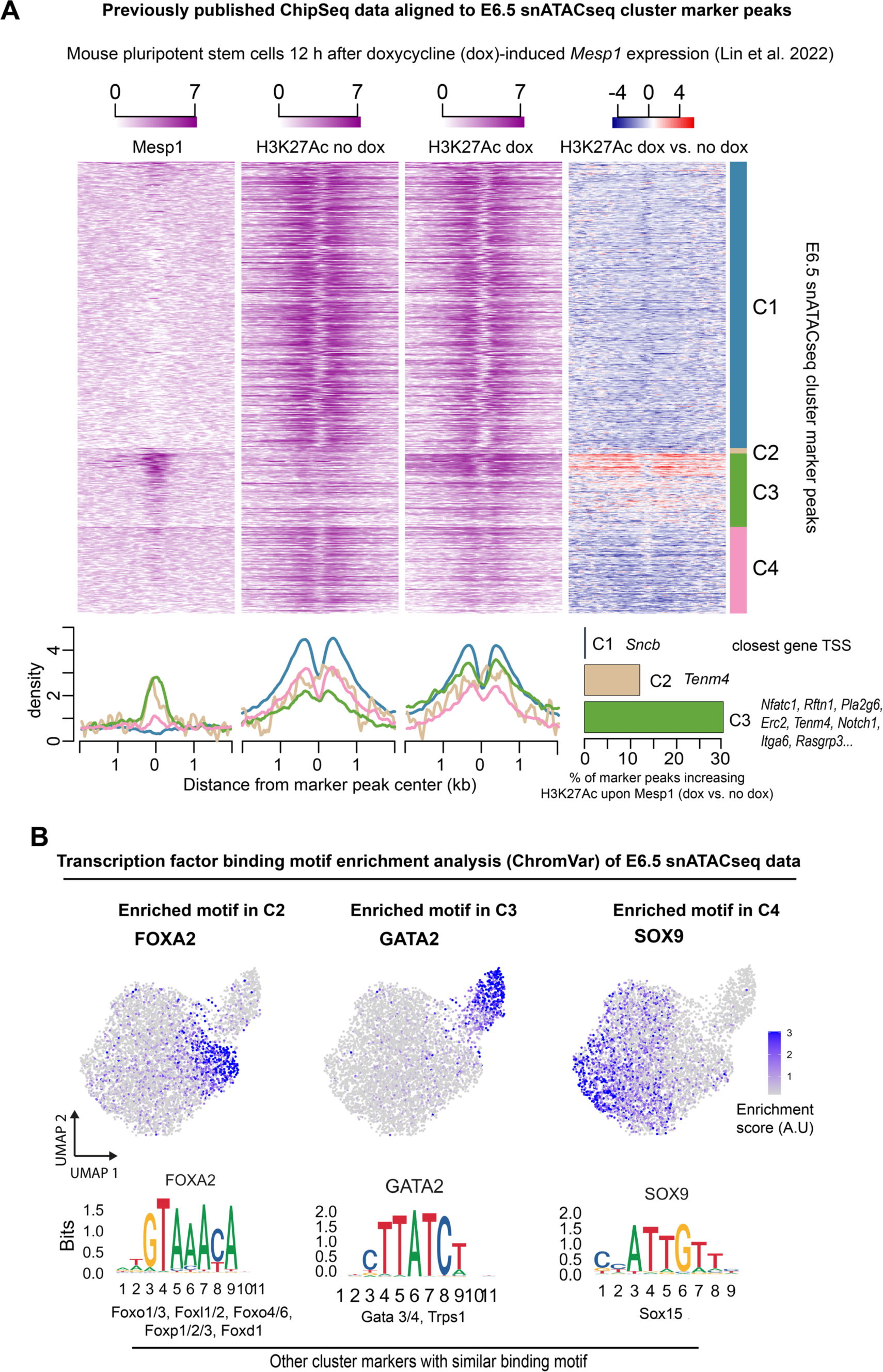
H3K27 acetylation around snATACseq E6.5 marker peaks in embryonic stem cells before and after Mesp1 expression. (A) Heatmap (stripplot) showing log2 normalized Chip-seq signal aligned on the E6.5 epiblast cluster marker peak regions. A fourth column shows the change caused by *Mesp1* induction in red-blue scale. Each row shows the ChipSeq signal around a marker peak. Data was obtained from ^27^, where mouse pluripotent stem cells 12 h after doxycycline (dox)-induced Mesp1 expression were analysed. The signal distribution for each cluster is summarized at the bottom. (B) Top transcription factor motifs enriched in the different E6.5 epiblast clusters.

To assess the degree of progress of these clusters towards gastrulation, we inferred the RNA expression of four key gastrulation transcription factors (*Fgf8*, *T*, *Eomes*, and *Mesp1*) by examining the accessibility of their coding sequences and promoter regions (**Fig. S2A**). *Fgf8*, *T* and *Eomes*, the earliest markers of prospective gastrulating cells in the epiblast (**Fig. S1_1**) showed sparse predicted expression in the ATACseq epiblast clusters, with cluster 2 and 3 showing higher frequency of *Eomes* expressing cells (**Fig. S2B**). *Mesp1* is the latest of the four genes to be expressed, being activated in the primitive streak at gastrulation onset (**Fig. S1_1**). None of the ATACseq clusters showed inferred *Mesp1* expression, suggesting that the epigenetic priming reported here precedes *Mesp1* expression onset (**Fig. S2B**). Overall, the accessibility of peaks associated with lineage-specific genes (**Fig. 1E; S2C**) and with DNA binding motifs of transcription factors involved in lineage specification (**Fig. 2B**), suggests that Cluster 2 contains precursors fated to endoderm, Cluster 3 contains mesodermal precursors ready for recruitment to the primitive streak and Cluster 4 cells that will remain as ectoderm (**Fig. S2D**). Clusters 0 and 1 did not show a clear epigenetic pattern and likely represent cells at an earlier point of progress towards differentiation.

These results show that the pre-gastrulation epiblast at E6.5 shows chromatin states related to different future cell lineages, with the strongest bias towards signatures of endothelial/endocardial fates. In contrast to the early appearance of the epigenetic marks found here, the first signs of endothelial differentiation in the embryo appear around E7.25-E7.5, coinciding with the onset of specific marker expression, such as the ETS Transcription Factor ERG^29^ (**Fig. 1G-H**). To understand how and when these epigenetic signatures translate into lineage specification through gastrulation, here we focused on the embryonic endothelial cell lineage.

### Specification of the endothelial versus non-endothelial mesodermal lineages

The strong endothelial chromatin signature in the epiblast, together with the absence of any bias towards other mesodermal fates, including cardiomyocytes, suggest independent specification of the endothelial lineage ahead of, and aside from other mesodermal lineages. The earliest endothelial cells to appear in the mammalian embryo belong to the first organ to develop; the primitive heart tube, which contains myocardium –composed of cardiomyocytes– and endocardium –composed of cardiac-specific endothelial cells–. Genetic lineage tracing, clonal analyses and stem cell experiments have supported the existence of early bipotential cardiac-specific progenitors responsible for generating both cardiomyocytes and endocardium^3,4,12,26,30–34;^ however, the potential of these progenitors to generate mesodermal lineages outside the heart was not investigated. Here, to describe the full set of lineage relationships of early embryonic endothelial/cardiomyocyte progenitors, we conducted a random, lineage-unrestricted clonal analysis using a tamoxifen-inducible ubiquitous driver –*RNApol2Cre–* and a two-reporter strategy^17,35–38^ (Methods). We adjusted the tamoxifen dose to target single cells during the ∼E6.25 to E6.75 stages and analysed the contribution of their progenies in whole embryos at E8.0-E8.25 (**Fig. 3A**, **B**). From 737 embryos generated, we focused on 44 showing fluorescence in the cardiac region containing a total of 46 labelled cell clusters. According to the two-reporter strategy statistics, 94.5% of monocolor cell clusters were expected to be derived from single cells (**Fig. S3_1H-J**, Methods). We analysed the contribution of each cluster to different embryonic compartments by immunostaining for sarcomeric Myosin heavy chain (MF20) and ERG, followed by confocal imaging to identify cardiomyocytes and endothelial cells (**Fig. 3C**). To quantify lineage relationships, we calculated the Jaccard similarity score for each possible combination (**Fig. 3D**). This revealed a weak lineage relationship between cardiomyocytes and endocardial cells. Instead, cardiomyocytes share greater lineage relationship with undifferentiated splanchnic mesoderm, while endocardial cells are more related to other endothelial cells in the embryo than to cardiomyocytes or any other mesoderm (**Fig. 3D**, **Fig. S3K, L**, supp. data 3).

**Figure 3:**
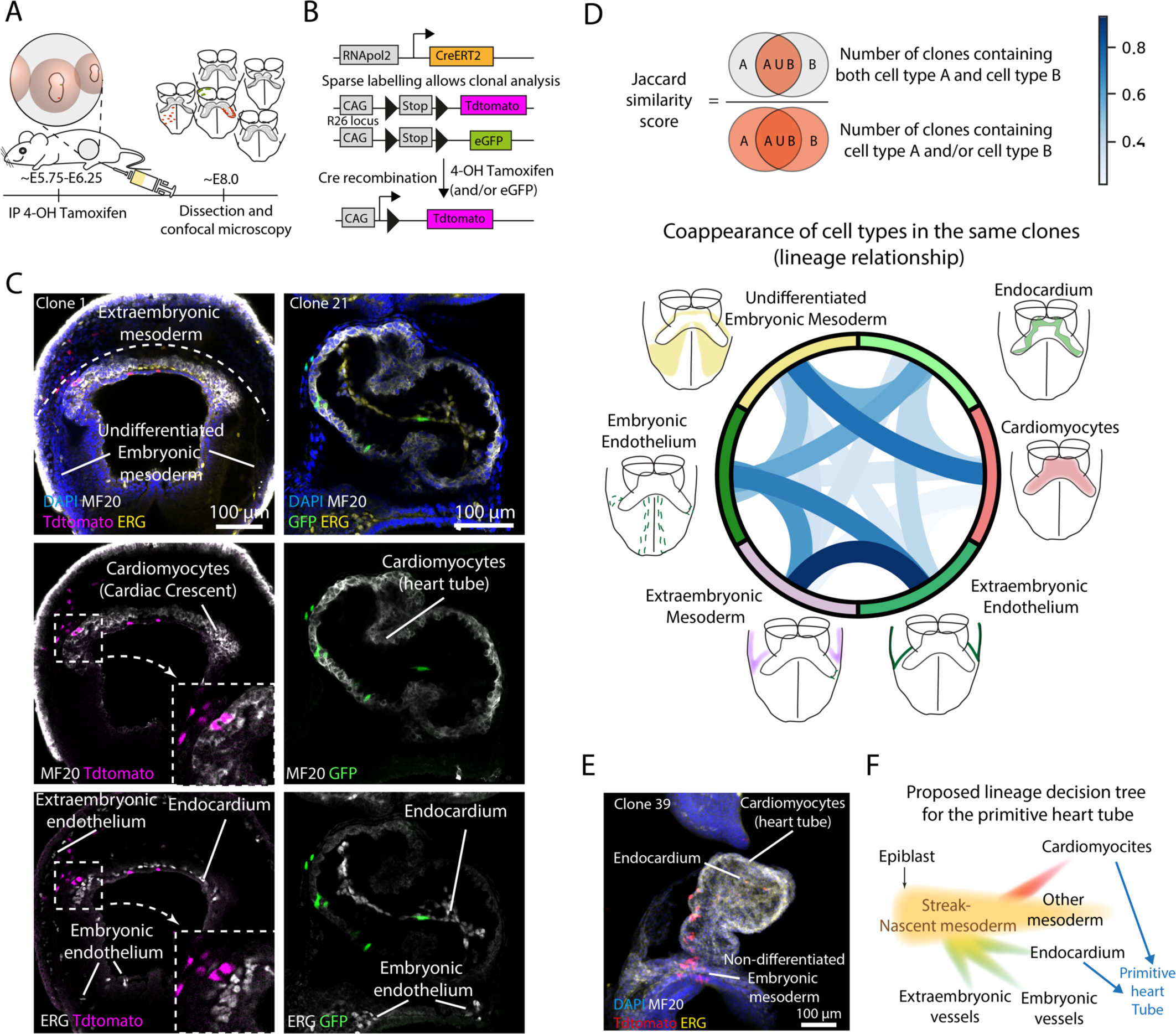
Retrospective clonal analysis of the primitive heart tube reveals independent specification of endothelial and cardiomyocyte lineages in the heart. (A) Clonal analysis strategy. (B) CreERT2 is expressed from the ubiquitous *RERT* allele^38^. As reporters of Cre activity, two *Rosa26* alleles, each driving the expression of Tdtomato or GFP, respectively. (C) Whole mount confocal optical planes showing the contribution of clones to mesodermal locations. MF20 and ERG immunostainings were used to distinguish cardiomyocytes and endothelial cells, respectively.(D) Chord diagram showing the Jaccard similarity score between the different cell types studied (n = 44 embryos). (E) Example of an unspecified clone that contributes to CM, EC, and cells outside the primitive heart tube. (F) Proposed lineage decision tree of the primitive heart tube from multipotent mesodermal progenitors, rather than bipotent cardiac-specific progenitors.

To investigate the timing of cardiomyocyte and endocardial cell specification, we estimated the embryonic stage at which recombination occurred. The induction time of each clone was estimated by calculating the time required to generate its number of cells according to the reported average cell division rate during the embryonic stages E6.5-E8.5^9,39^ and subtracting this time from the actual stage at dissection (**Fig. S3_1A** and **B**, Methods, supp. data 4). Our estimation aligned with the reported pharmacodynamics of 4-OH tamoxifen in mouse blood, which peaks around 12 hours after injection (**Fig. S3C,** supp. data 5). In addition, the scoring of bilateral clones, which only result from progenies labelled before mesoderm ingression^40^, identified the timing of primitive streak ingression for cardiac progenitors around E6.75 (**Fig. S3D-F**). The chronological ordering of clones revealed that inductions resulting in progenitors that produced both cardiomyocytes and endocardial cells occurred before E7.0 and contributed to other mesoderm regions outside the primitive heart tube in most cases (17 out of 19, **Fig. S3G**).

Next, we performed prospective clonal analysis by TAT–Cre microinjection^41^, which allowed us to recombine single cells at custom stages and embryonic locations, and cultured the embryos to analyse the resulting clones in the heart (**Fig. 4**. suppl. data 6). We obtained clones contributing to both cardiomyocytes and endocardium at high frequency (5/7) when injecting pre-streak embryos and at lower frequency (1/4) when injecting early-streak embryos, whereas later injections always labelled separate lineages (**Fig. 4**). These results show that the endothelial and cardiomyocyte lineages are already independent at the time of their ingression in the primitive streak (∼mid-streak). Clones that showed mixed progenies again also contained cells outside of the primitive heart tube, consistently with the low frequency of cardiomyocyte-endocardial exclusive clones in the tamoxifen-induced samples. Together, snATACseq and clonal analyses show that endothelial cells are independently specified from other mesodermal cells –including cardiomyocytes–, and subsequently are recruited together with cardiomyocyte precursors to form the heart (**Fig. 3F**).

**Figure 4:**
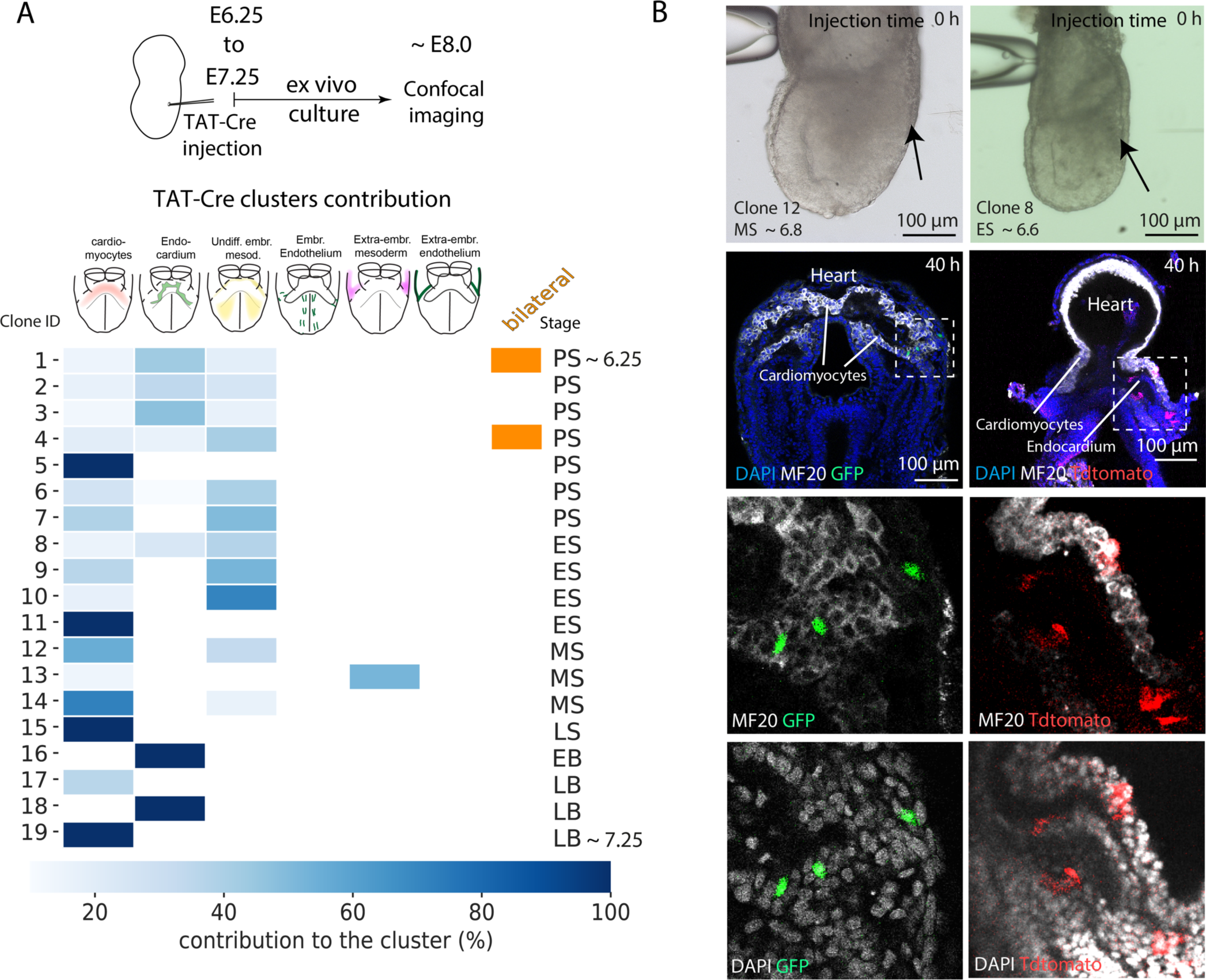
TAT–Cre microinjection for prospective clonal analysis of primitive heart tube progenitors. (A) Experimental setup and contribution of TAT–Cre–induced clones. (X axis – clone, Y-axis – location. n = 19 clusters, 18 embryos, 8 litters). (B) Two embryos being injected, and the resulting clone (confocal plane). In the left embryo, zoom–ins reveal that recombined cells are located both inside and outside the primitive heart tube, stained for the myocyte marker MF20. On the right, an unspecified progenitor contributed to cardiomyocytes, endocardium, and undifferentiated mesoderm.

### Spatiotemporal mapping of cardiac precursors

Our results indicate an important temporal gap between the specification and differentiation of the endothelial cells, as detected by specific marker gene expression. To understand the emergence of endothelial differentiation during this gap, we used live imaging to track cardiac progenitors from ∼E6.75 to their differentiation in the cardiac crescent at ∼E8.0. Adapting our previous protocol^42,43^, we conducted time-lapse 3D imaging using two-photon microscopy and random cell lineage tracing using *CreERT2* and *Tdtomato^flox/flox^* reporter alleles^36,38^ (**Fig. S5A-C**). Additionally, we used a *CBF1-Venus* allele to identify endothelial cells at later timepoints^44^ (**Fig. S5D, supp.** data 7). First, based on tissue morphology we identified the cardiac crescent region at the final timepoints of the videos. Second, we distinguished cardiomyocytes as rounded cells forming a chamber and endocardial cells as Venus-positive elongated cells within the chamber lumen. Finally, we manually tracked back cardiomyocytes and endocardial cells to the beginning of the videos using the MaMut ImageJ plugin^45^ (**Fig. S5C-E**). We applied this strategy to analyse two *CreERT2*; *Tdtomato^flox/flox^*; *CBF1-Venus* embryos, tracing Tomato+ cells and a *H2B:miRFP703* embryo with ubiquitous nuclei fluorescence^40,46^ (videos 1-3).

By detecting cell divisions during tracking, we reconstructed the lineages of individual progenitors in the nascent mesoderm surrounding the primitive streak (**Fig. 5A-B**, video 6). After excluding lost tracks, these progenitors were linked to 146 descendant cells at the end of the time-lapse sequences (**Fig. 5C-D**). Among the progenies we studied, we did not find single cells yielding both cardiomyocytes and endocardial cells. In total, we identified 15 cardiomyocyte and 16 endothelial progenitors, which divided on average every 7.4 h and 7.6 h, respectively, during the observation period (**Fig. 5SF**). The equivalent division rate suggests that the proportion of initial progenitors specified is the same as the proportion of cardiomyocytes and endothelial cells found in the primitive heart tube, approximately four to one (**Fig. 5SG**). In line with the clonal analyses, cell tracking shows that the endothelial lineage is already segregated at the beginning of the time-lapse study.

**Figure 5:**
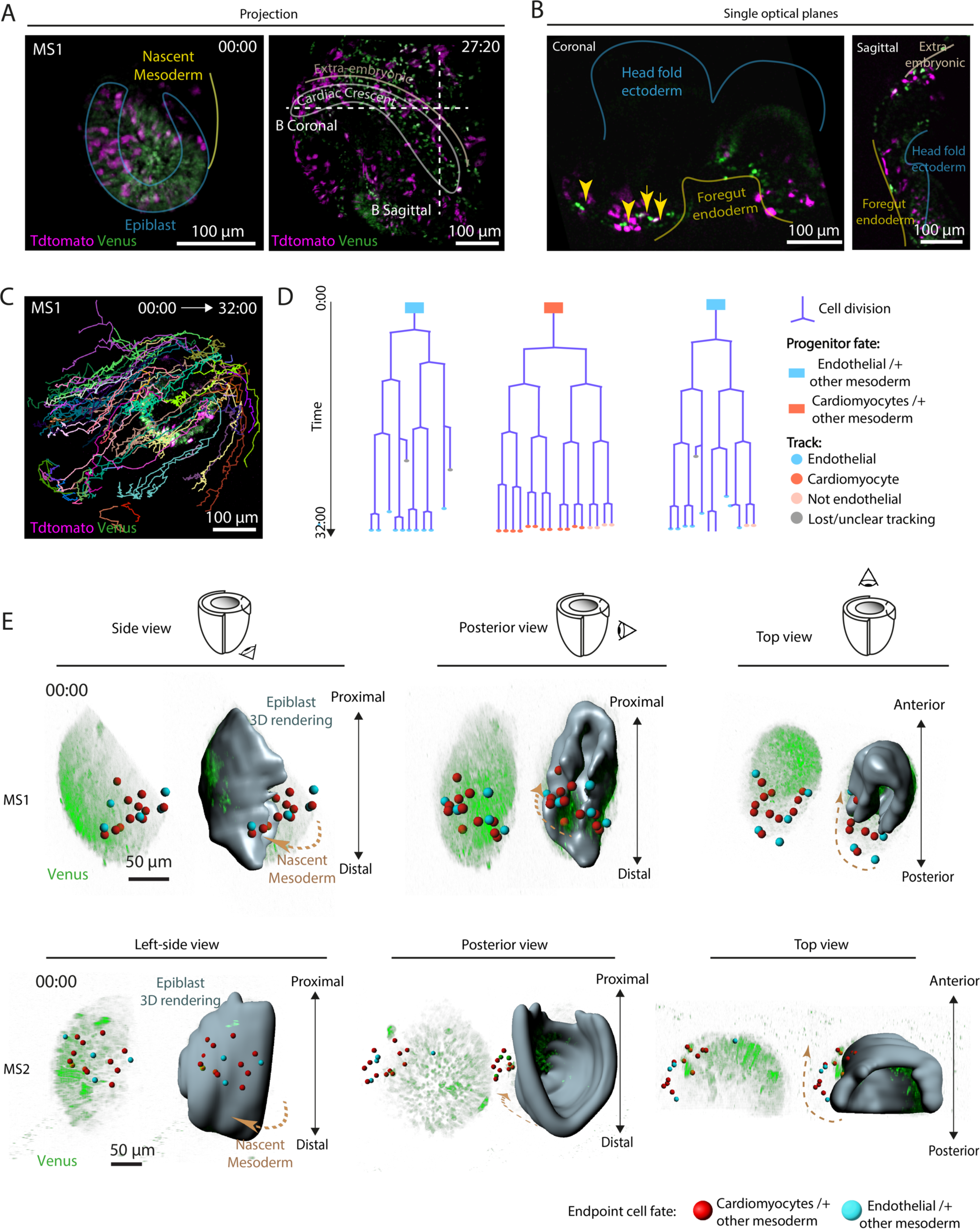
Two–photon time–lapse microscopy for cardiac progenitor’s cell tracking and lineage reconstruction. (A) Example of live imaging tracked embryo. Intensity projections at initial and final time points. (B), Coronal and sagittal optical planes of the same embryo, highlighting main morphological features of the embryo. Yellow arrows indicate endothelial cells expressing the *CBF1:Venus* transgene, which allows identification of endothelial cells. (C) Cell tracks from the beginning to the end of the tracking as displayed in MaMuT Viewer. (D) Examples of reconstructed lineage trees from the earliest progenitor to all cell descendants. Endpoint cell fate of progenitors is depicted as a coloured square; progeny cell type as coloured circles. Each branch bifurcation represents a cell division event. (n = 31 and 146 cells at initial and final time points, respectively, from 3 embryos). (E) Different views of raw data and 3D epiblast volume rendering two different embryos. Red dots indicate the initial position of non– endothelial progenitors, while blue dots indicate the position of endothelial progenitors.

In addition, these results provided spatiotemporal information about cardiac progenitors. We then used this information to map the positions of the progenitors at their exit from the primitive streak, which revealed no spatial segregation of endothelial and non-endothelial cardiac progenitors (**Fig. 5E**). This suggests that, despite being already segregated, cardiomyocyte and endocardial cell progenitors arise simultaneously from the same primitive streak region and are therefore exposed to equivalent signalling environments at this point.

### Differential migration behaviour in endothelial progenitors prior to mesoderm epithelization

Leveraging the 3D + time trajectories obtained in *H2B:miRFP703* cell tracks, we further explored the evolving migratory behaviour of endothelial and non-endothelial cardiac progenitors throughout gastrulation and heart tube assembly, in order to identify the onset of cardiac progenitor differentiation (**Fig. 6A**). We assessed 15 kinetic parameters in the tracks of both endothelial and non-endothelial progenitors, including speed, straightness, and distance to the endoderm, which revealed a gradual divergence between the two progenitor types (Methods, **Fig. S6A**). Using a linear mixed-effects model, we compared both groups across five temporal windows on each of the parameters, revealing that the first differences for some of the parameters appear in the second time window, E7.0-E7.25 (**Fig. S6B,** supp. data 8). During this time period, prospective endothelial cells become faster, more exploratory and move to positions closer to the endoderm (**Fig. S6A, B**).

**Figure 6:**
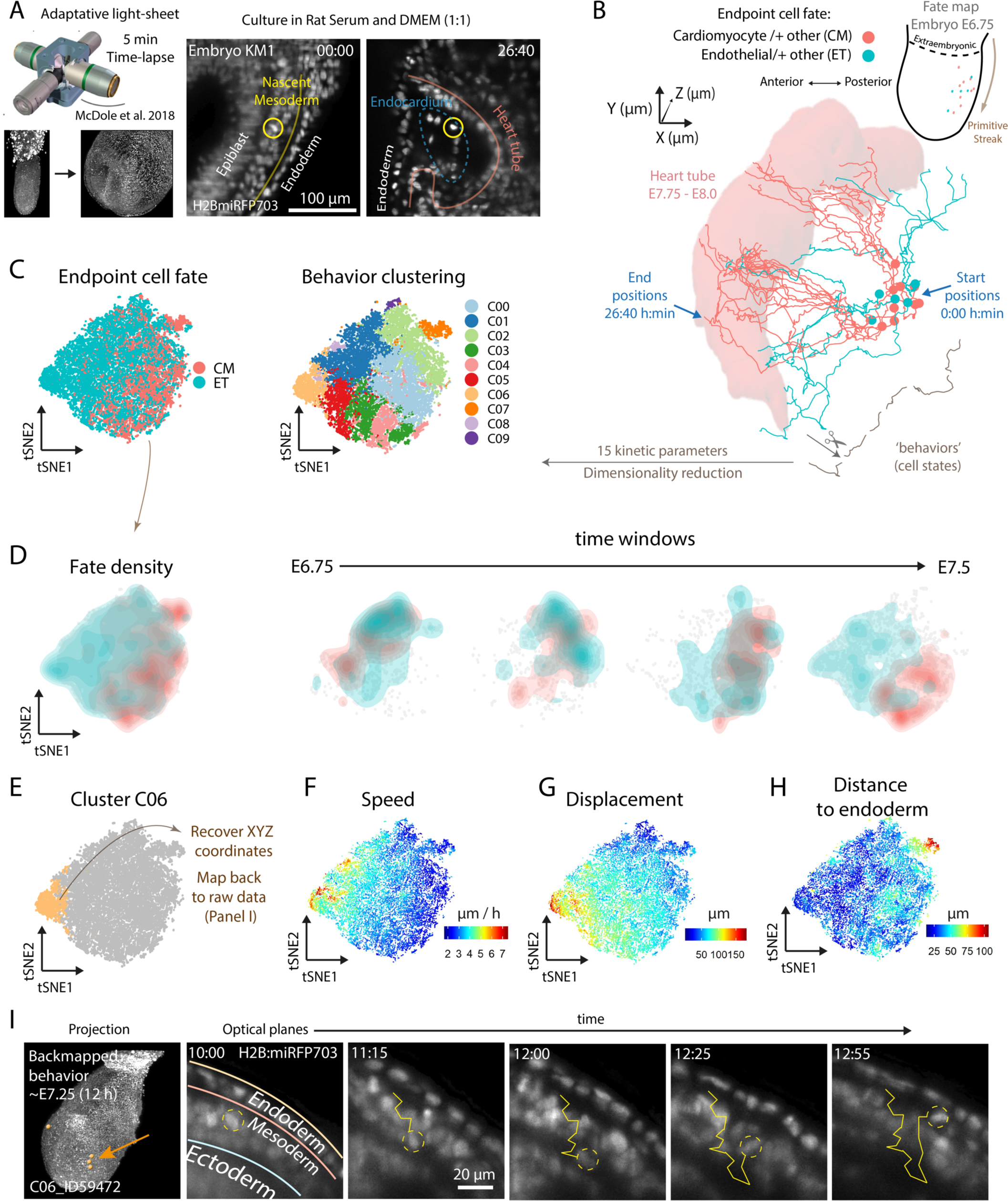
Live imaging reveals differential migration behaviour in endothelial progenitors prior to mesoderm epithelization. (A) Right, Imaging setup in^52^; left, Initial and final time points of an endocardial progenitor track in a *miRFP703*. (B) 3D representation of KM1 embryo cell tracks. The XYZ coordinates of endothelial (green) and non–endothelial (red) heart tube progenitors are lined. The initial positions are shown as spheres, and the location of the heart tube at the final time point is shown as a red rendering for clarity. (C) t-SNE (T-distributed Stochastic Neighbour Embedding) representations of cell migration states colour by endpoint cell fate (left) and cluster (right) (17,170 cell states from 173 cells imaged over 26 h). (D) t-SNE density plots of endpoint cell fate divided by 4 periods. (E) Cluster C06 in t-SNE. (F-H) Different measured migration parameters in t-SNE. (I) Back-mapping of cluster 06 cell states to the raw data. Timepoints of a representative track are shown.

We next aimed to analyse all parameters collectively and unbiasedly along developmental progression, providing a comprehensive view of cell behaviour. However, cells within a shared pool of progenitors may traverse the differentiation trajectory at slightly varied embryonic stages. For instance, two endocardium progenitors undergoing gastrulation at different times may not be concurrently exposed to inductive signals, leading to potential differentiation delays, yet they belong to the same population. To classify cell states regardless of embryo stage, we employed a methodology akin to that used for classifying immune cell behaviours^47^. First, we fragmented cell tracks to represent distinct cell states –similar to clips that play moments in the trajectory of a moving object (Methods, **Fig. 6B**). Then, using the 15 kinetic parameters, we plotted cell states on a t-distributed stochastic neighbour embedding (t-SNE, **Fig. 6C**). Clustering identified ten main behavioural groups with differential enrichment in endothelial or cardiomyocyte- fated tracks (**Fig. 6C, Fig. S6C-E)**. Given that the analysis is time-resolved, we could determine that in earlier time points, endothelial and non-endothelial progenitors exhibited similar behaviours, but from E7.25 onward, they segregated, indicating the emergence of behavioural differences (**Fig. 6D**).

Mapping the cell states contributing to a specific cluster back to the raw data allowed us to scrutinize their location and migration signature (**Fig. 6E-H**). The earliest behaviours prevalent in endothelial progenitors were characterized by high speed and high exploratory activity (C06, **Fig. 6E-I**, **Fig.S6D**, **E**, video 5), contrasting with the more parsimonious migration of non-endothelial progenitors (C04, video 6). The behaviour of endothelial precursors at this stage denotes a clear divergence from the main stream of still migrating global population of mesodermal precursors (video 5). The early segregation of endothelial behaviours precedes mesoderm epithelization and coelomic cavity formation, challenging the current conception of endocardial cells first differentiating by delamination from the pre-cardiac epithelialized mesoderm^12^. Instead, our live imaging data show that endothelial progenitors start differentiation during mesoderm migration and before mesoderm epithelialization. These observations support the early differentiation of the endothelial lineage from the rest of mesodermal derivatives shortly after gastrulation.

## Discussion

Here we showed that chromatin accessibility indicates cell lineage priming in the mammalian epiblast. Chromatin signatures in the E6.5 epiblast suggest pre-specification of cell populations to either the mesodermal, endodermal or ectodermal fates. Strikingly, we found that the strongest lineage bias in the epiblast corresponds to the endothelial fate, which reveals a dedicated program for the early specification of the endothelial lineage in the mammalian epiblast. This nuanced epigenetic landscape suggests a new schedule for lineage specification in the mammalian epiblast. Contrary to the idea of a bipotential common cardiac progenitor, clonal analysis and live imaging revealed early progenitors contributing to multiple lineages and their rapid transition to endothelial specification without progressive bifurcations. Furthermore, endocardial cells are more related to other endothelial cells than to cardiomyocytes, which reinforces the idea of independent specification of the endothelial cell lineage, followed by recruitment of cardiomyocyte and endothelial precursors to form the myocardium and endocardium.

Using live imaging, we tracked cardiomyocytes and endothelial cells back to their initial positions in the nascent mesoderm, revealing no spatial segregation. This suggests that these progenitors arise from intermingled positions in the primitive streak, ruling out differential exposure to morphogens in the primitive streak as a likely mechanism for their specification^8,48^. One possibility is that local cell interactions or random mechanisms pattern the emergence of the endothelial lineage in the epiblast or in the primitive streak; however, we cannot exclude regional specification in the epiblast followed by subsequent co-recruitment to the same region of the primitive streak.

Analysing the migration behaviour of fate-assigned progenitors revealed distinctive endothelial behaviours preceding splanchnic mesoderm epithelialization and coelomic cavity formation. This finding resets the onset of endothelial cell differentiation to earlier times than previously noticed using classic approaches, and challenges the established model of endocardium development, where endocardial progenitors delaminate from the cardiac mesoderm post-epithelialization^12^.

Together, our findings reshape our understanding of early embryonic development, highlighting the role of epigenetic priming in pluripotent epiblast cells. This priming may prepare endothelial progenitors for a swift differentiation following gastrulation, while allowing them to transiently keep their differentiation schedule dormant for ensuring their proper delamination and migration through the primitive streak. In the primitive streak, and upon expression of mesodermal transcription factors^24^, they would rapidly transition to an active state, allowing them to respond to inductive signals from the anterior visceral endoderm at E7.25, ensuring the timely formation of the primitive heart tube and embryonic vasculature. The silent epigenetic priming of endothelial precursors in the epiblast therefore would allow them to remain on-hold for differentiation while allowing the execution of the gastrulation program, followed by early differentiation onset. In contrast to the early differentiation of endothelial cells reported here, cardiomyocyte differentiation only starts once cells arrive to the cardiogenic region and epithelialize^42,49^, and therefore, early priming of the cardiomyocyte lineage would be dispensable.

## Supplementary files

<u>Video 1</u>. Embryo MS1. Live imaging *CBF1:Venus RERT Tdtomato* embryo.

<u>Video 2</u>. Embryo MS2. Live imaging *CBF1:Venus RERT Tdtomato* embryo.

<u>Video 3</u>. Embryo KM1. Live imaging *H2B:miRFP703* embryo.

<u>Video 4</u>. Example of endothelial progenitor track with annotated cell layers. Embryo KM1. Live imaging *H2B:miRFP703* embryo.

<u>Video 5</u>. 4D tracking behavioural analysis. Example of cluster 05 cell behavior. Mesodermal, endothelial progenitor cell.

<u>Video 6</u>. 4D tracking behavioural analysis. Example of cluster 04 cell behavior. Mesodermal, non-endothelial progenitor cell.

<u>Supplementary data 1</u>. snATACseq differentially accessible peaks for the epiblast clusters with closest genes and genes sharing the same regulatory region. (Data for Fig. 1E).

<u>Supplementary data 2</u>. snATACseq predicted transcription factor motif activity enriched in different epiblast clusters (Data for Fig. 2B).

<u>Supplementary data 3</u>. Retrospective clonal analysis metadata. Sheet 1: Contribution of tamoxifen-induced clones to different mesodermal compartments, number of cells, estimated recombination stage and estimated delay between tamoxifen injection and recombination on each clone (Data for Fig.S3). Sheet 2: Coappearence matrix of cell types (Jaccard similarity score) (Data for Fig.3D). Sheet 3: Retrospective clonal analysis embryo stage at dissection and fluorescence. Sheet 4: Retrospective clonal analysis litter metadata. Tamoxifen dose, time of injection, mice weight and time of dissection.

<u>Supplementary data 4</u>. Prospective clonal analysis metadata. Contribution of TAT-Cre induced clones to different mesodermal compartments, number of cells, time and region of injection and time of *ex vivo* culture (Data for Fig4).

<u>Supplementary data 5</u>. Live imaging. Sheet 1: Quantification of *CBF1:Venus* + cells in the endothelium marked by CD31 staining (Data for FigS5D). Sheet 2: Division time in tracked cells (Data for FigS5F). Sheet 3: Ratio MF20 (cardiomyocytes, CM) over ERG (Endothelial cells, EC) (Data for FigS5G).

<u>Supplementary data 6</u>. 4D behavioural analysis. Univariate statistical analysis with a linear mixed-effects model to compare track parameters along 5 different time windows.

## Acknowledgements

We express gratitude to Kenzo Ivanovitch for his guidance in live imaging techniques and mentoring M.S. at the outset of this research. We thank members of the Microscopy and Dynamic Imaging, Transgenesis, and Animal Facility CNIC units for excellent support. We thank “la Caixa” Foundation (ID 100010434) for the fellowship that supported M.S. stipend (LCF/BQ/DE18/11670014). We also thank The Company of Biologists for the travelling fellowship that made possible M.S. stay at Janelia Research Institute with K.M. and P. K. (DEVTF181145). This work was funded by grants PGC2018-096486-B-I00 and PID2022-140058NB-C31 from the Agencia Estatal de Investigación to M.T.; European Commission H2020 Program grant SC1-BHC-07-2019. Ref. 874764 “REANIMA” to M.T., Comunidad de Madrid grant P2022/BMD-7245 CARDIOBOOST-CM to M.T. The CNIC Unit of Microscopy and Dynamic Imaging is supported by FEDER ‘Una manera de hacer Europa’ (ReDIB ICTS infrastructure TRIMA@CNIC, MCIN). The CNIC is supported by the Ministerio de Ciencia e Innovación and the Pro CNIC Foundation and is a Severo Ochoa Center of Excellence (CEX2020-001041-S).

## Author contributions

Conceptualization: M.S., M.T.; Methodology: M.S., J.N.D, J.d.D.H.; Software: M.S., D.J.C; Validation: M.S., J.N.D.; Formal analysis: M.S., J.N.D.; Investigation: M.S., J.N.D.; Data curation: M.S., J.N.D.; Writing - original draft: M.S.; Writing - review & editing: M.S., M.T.; Supervision: M.T.; Project administration: M.T.; Funding acquisition: M.T.

## Code availability

Code to reproduce the analysis in this paper will be available at: https://github.com/MiquelSendra/EpiLineagePriming

## Data availability

Ready to use IGV browser session with bigwig tracks is available here https://drive.google.com/drive/folders/1NuODkvpF9iz5dnkQePkSPKYQPdiyvFJV?usp=sharing

Just copy the whole content of the folder in your computer and load the igv_session.xml file in IGV. The paths are relative so it will find all bigwig tracks within the same folder.

Processed snATACseq data are available as a Seurat object here: https://drive.google.com/drive/folders/1HByJyAdu5Ve2J2UUydVmcXMyKR97IU8E?usp=sharing

Processed 4D tracking behavioural data are available as a Seurat object here: https://drive.google.com/drive/folders/1-pqBVsV0YOroG6qnKgLHAZuP_e2gmKJJ?usp=drive_link

## Materials and Methods

### Mouse strains

Animals were handled in accordance with CNIC Ethics Committee, Spanish laws and the EU Directive 2010/63/EU for the use of animals in research. All mouse experiments were approved by the CNIC and Universidad Autónoma de Madrid Committees for “Ética y Bienestar Animal” and the area of “Protección Animal” of the Community of Madrid with reference PROEX 220/15. For this study, mice were maintained on mixed C57Bl/6 or CD1 background. We used the following mouse lines (Table III.1, which were genotyped by PCR following the original study protocols. Male and female mice of more than 8 weeks of age were used for mating.

| Mouse line | Reference |
| --- | --- |
| Polr2a-CreERT2 (RERT) | Guerra et al. (2003) |
| ROSA26CAG-TdTomato | Madisen et al. (2010) |
| ROSA26CAG-EGFP | Sousa et al. (2009) |
| Tg(CBF:H2BVenus,+) | Nowotschin et al. (2013) |
| Tg(H2B:miRFP703,+) | Gu et al. (2018) |

### Embryo retrieval

Embryos were staged considering 12:00 on the midday of the vaginal plug as embryonic day (E) 0.5. Females were sacrificed by cervical dislocation. The abdominal cavity of sacrificed females was opened to expose the uterus. The uterus was then placed in ice cold PBS for fixed analysis or in 37◦C dissection media for experiments requiring embryo culture (see Methods). After opening the muscle layer and the decidual layer, the embryos were extracted, dissected and finally fixed in paraformaldehyde (PFA, Merck) 2% in PBS overnight at 4◦C or placed in pre–equilibrated culture medium.

### Whole mount embryo immunostaining

After fixing embryos in 2% PFA in PBS, immunofluorescence was performed as follows: After three washes with PBS, the embryos were permeabilized with a 0.3% Triton X-100 in PBS solution for 30 minutes at room temperature. Blocking was performed with Bovine Serum Albumine (BSA, Thermo Fisher) 0.5% in PBS for at least 3 hours at 4◦C. The primary antibodies were then incubated overnight. We used the following primary antibodies: anti–CD31 (553370 BD Pharmingen clone MEC 13.3), anti– M20 (1:100; Anti–MF–20–mouse Hybridoma bank), anti–ERG (1:500; Anti–ERG antibody Rabbit [EPR3863]-–ChiP Grade, Abcam Ab110639). Primary antibody washing was carried out in a 0.1% Triton X–100 in PBS solution for at least 5 hours at 4◦C. Secondary antibody incubation was carried out overnight at 4◦C. We used the following secondary antibodies: Alexa Fluor 647 goat anti–mouse (1:500; Life technologies A31571) and Alexa Fluor 594 goat anti–rabbit (1:500; Life technologies A11037). For anti–6xHis-tag staining, embryos were washed for 2 days at 4◦C and then incubated for 5 minutes with TSA Cyanine 5 at room temperature (NEL705A001, Akoya, biosciences). All embryos were nuclei stained with DAPI 1:1000 diluted in PBS. Embryos were clarified in crescent dilution of glycerol in PBS (25%, 50% and 75%) until analysis was performed by confocal microscopy.

### Confocal microscopy of fixed embryos

Whole embryos were mounted on 35 mm plates with a 14 mm diameter glass coverslip (Mattek, P35G-1.5-14-C) and imaged on a Leica TCS SP5 confocal microscope using 405, 488, 561, 633 nm wavelengths and 10x/0.4 dry and 20x/0.75 glycerol objectives or on a Leica TCS SP8 confocal microscope using spectral wavelength lasers and nd 20x/0.75 glycerol objectives. A 3D stack was obtained by imaging optical sections every 3 or 5 µm.

For quantification of the Notch reporter expression (CBF1:H2BVenus, +) in the endothelium, we used immunostaining of the CD31 marker. Following confocal imaging, CBF:H2BVenus positive and negative cells were counted within the CD31 positive and negative domains using ImageJ Cell Counter plugin, which output was plotted and statistically analyzed using chi–square test to compare the proportions of positive cells in both domains.

### Retrospective clonal analysis

For retrospective clonal analysis, we used mouse embryos carrying the inducer *Polr2a– CreERT2 (RERT)* and both the reporters *ROSA26CAG–TdTomato* (R26RtdTomato) and *ROSA26CAG–EGFP* (R26REGFP) in trans-heterozygosis. These genotypes were generated upon breeding mice that have the inducer and one of the reporter alleles in double homozygosis with mice that have the second reporter allele in homozygosis. Random Cre–mediated recombination was triggered with 4–hydroxy–tamoxifen dissolved in corn oil. A single dose of 4–hydroxy–tamoxifen was injected intraperitoneally into pregnant females at E5.75 or E6.25 days of gestation. The embryos were dissected, fixed and analyzed at E8.0–E8.5 as described in the following sections.

### Tamoxifen preparation

For induction of the RERT line, 10 mg of 4–hydroxy Tamoxifen (Sigma) was dissolved in 1 ml of absolute ethanol and 9 ml of corn oil (Sigma) for a final concentration of 1 mg/ml. The stock solution was then sonicated for 40 minutes on ice to prevent overheating. The solution was aliquoted and stored at 4◦C for up to 4 weeks, and re–sonicated before being administered to mice.

### Cluster cell counting

Once acquired, the images were opened as optical plane stacks and saved in .tiff format. The contribution of the clusters to each anatomical location was evaluated by counting DAPI nuclei within Tdtomato+ or GFP+ cells. Anatomical locations were identified using morphological features (Kaufman and Navaratnam, 1981) in the DAPI channel. Additionally, MF20 and ERG immunostaining signal identified cardiomyocytes and endothelial cells. Two overlapping groups of cells (Tomato and GFP cells) were annotated as “bicolor clusters”.

The polyclonality of monocolour cell clusters in the embryo collection was estimated using the frequency of bicolour clusters as previously described^2,35^. This method is based in the fact that the frequency of bicolour events in a collection of samples is directly proportional to its polyclonality (Figure III.1A). This allows calculating the probability of finding clusters labeled with one reporter (monocolour clusters) that originate from multiple progenitors (polyclonal) using the following formula (Figure III.1F). We reported previously the relative recombination of GFP and Tdtomato reporter^41^. Briefly, we first estimated the relative Tomato and GFP recombination frequency: *RERT+/-;ROSA26RCAG–TdTomato+/+* mice were crossed with *ROSA26RCAG–GFP+/+* mice. Reporter recom- bination was induced by administering 0.04 mg/g of 4–OH tamoxifen intraperitoneally to pregnant females on day E7. A day later, the embryos were harvested and the relative efficiency of recombination was calculated by manually counting GFP and TdTomato cells over total DAPI nuclei in confocal optical sections using the ImageJ Cell Counter plugin. We found that Tdtomato recombined 1.97 times as often as GFP.

### Clonal probability

The frequency of mono–color polyclonal clusters can then be estimated as a function of the frequencies of bicolor clusters and of mono–color clusters, which is biased towards the production of Tdtomato clusters, with a calculated frequency of recombination of 1.76% for Tdtomato and 0.87% for GFP (that is, Tdtomato recombines 2.02 times more often). Dismissing polyclonality levels above biclonality and assuming a stochastic distribution of clusters, the following functions apply for estimations:

- ▪ Frequency of bicolor clusters = frequency of Tdtomato (A) x frequency of GFP (B) × 2
- ▪ Frequency of polyclonal monocolor clusters= A2 + B2
- ▪ Then, the frequency of polyclonal monocolor clusters = Frequency of bicolor clusters 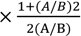

In our case, for the retrospective clonal analysis, 4.5% of the clusters analyzed (2/44) were bicolor. Applying these formulae, we calculated that the clusters in our collection had a 94.3 % chance of being actual clones.

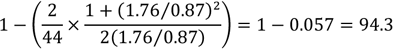

### TAT–Cre prospective clonal analysis

Mouse embryos at developmental stages E6.5 to E7.5 were dissected in a pre-equilibrated medium containing DMEM supplemented with 10% fetal bovine serum, 25 mM HEPES-NaOH (pH 7.2), penicillin, and streptomycin. Subsequently, the embryos were cultured under controlled conditions within a hypoxic chamber incubator at 37°C with 5% O2 and 7% CO2, using a culture medium comprising 50% Janvier Labs Rat Serum SPRAGUE DAWLEY RjHan SD male only and 50% DMEM FluoroBrite. For prospective clonal analysis tracing, embryos in the same developmental range were microinjected with TAT-Cre recombinase using specialized equipment and techniques. Specifically, microinjection needles were prepared with a 2µm gauge and inserted into the anterior side of the embryo until penetrating the endodermal layer, using specified pressure conditions. The embryos were handled and positioned carefully, ensuring that the anterior and posterior sides were oriented accordingly during the procedure to achieve successful microinjections.

In our prospective clonal analysis, we utilized mouse embryos that carried both the reporter genes ROSA26CAG–TdTomato (R26RtdTomato) and ROSA26CAG–EGFP (R26REGFP) in transheterozygosis. Following a process akin to the one used for retrospective clonal analysis, we fixed, imaged, and annotated fluorescent cells within anatomical regions. “Clusters” were defined as groups of cells (either Tomato or GFP) originating from a single TAT-Cre injection. We employed the same probability calculation method as in retrospective clonal analysis, using the two-reporter strategy as previously outlined in the literature^2,35^. Notably, out of 19 clusters analyzed, only one was bicolor, indicating that clusters in the TAT-Cre induced embryo collection had a high likelihood (93%) of being monoclonal.

### Embryo culture and live imaging of gastrulating mouse embryos

Live imaging procedures followed the protocol outlined in ^43^. brief, mouse embryos were carefully collected and dissected within a dissection medium comprised of DMEM supplemented with 10% fetal bovine serum, 25 mM HEPES-NaOH (pH 7.2), and penicillin-streptomycin (50 µg/ml each). For embryos spanning E6.5 to E7.5, culture conditions were established using a mix of 50% Janvier Labs Rat Serum Sprague Dawley RjHan SD (male only) and 50% DMEM FluoroBrite (Thermo Fisher Scientific, A1896701) with incubation at 37°C and a 7% CO2 concentration. Imaging was conducted on a Zeiss LSM780 platform, featuring a 20× objective lens (NA=1) and a MaiTai laser set at 980 nm for two-channel two-photon imaging. Fluorescence was detected with Non Descanned Detectors equipped with the filters cyan-yellow (BP450-500/BP520-560), green-red (BP500-520/BP570-610) and yellow-red (BP520-560/BP645-710). Zen software (Zeiss) facilitated data acquisition with an output power of 250 mW, pixel dwell time of 14.8 s, line averaging of two, and an image dimension of 610×610 µm (1024×1024 pixels).

### Cell tracking, lineage reconstruction from live imaging data

We used Leo Guignard’s lab rigid block–matching registration tools (<u>GitHub repository</u>) developed initially by Grégoire Malandin and Sebastien Ourselin ^40,53^ and later optimized by Leo Guignard for this project. Block–matching registration corrects translation and rotation in all of the planes. It does so by making blocks of the images and trying to match the intensities between one time point and the next. Subsequently, the blocks are made smaller until optimal matching is found. This corrects for embryo drift and sudden motion between one time point and the next, allowing for the quantification of cell tracking parameters and facilitating tracking itself.

To reconstruct lineages and assess the specification of early cardiac progenitors in our live imaging data, we tracked differentiated cardiomyocytes and endothelial cells located in the cardiac crescent or primitive heart tube back to their initial positions in the nascent mesoderm. Manual cell and lineage annotations were performed using the Fiji plugin Massive Multi-view Tracker (MaMuT)^45^. Cells that could not be reliably identified in the previous or following time points were discarded. Once a progenitor was successfully tracked from the beginning to the end of the video, the rest of its sisters were tracked on each division to reconstruct the full lineage. For some cases, sister cell tracks were lost due the cell falling out of frame, moving to an area with poor resolution or crowded with many cells. Next, MaMut output files (parsed .xml in a graph data structure) were processed using a custom python script. This script is available as a Jupyter notebook in our GitHub Repository. Briefly, we used Leo Guignard’s LineageTree Python library (<u>GitHub repository</u>) to retrieve cell lineages and XYZ coordinates from the .xml files to plot tracks in 3D and lineage representations.

### Signature scores on previously published Mouse Gastrulation Atlas scRNAseq data

R notebooks are available in our GitHub repository. Briefly, data was loaded using MouseGastrulationData package. Marker gene lists were imported from the markers tab in MouseGastrulationAtlas browser (https://marionilab.cruk.cam.ac.uk/MouseGastrulation2018/). Using UCell package^23^, we calculated signature scores for each cell type at E6.5, E6.75, E7.0 and E7.25 using the ScoreSignatures_UCell function. UMAP plots were generated to visualize the scores for each cell type, and a custom color palette was used to distinguish between different score levels.

### Nuclear isolation for snATACseq

The uterus was removed from 6-day pregnant dams at 9:00 am as previously described^43^. 5 females yielded a total of 46 embryos. We discarded embryos with signs of primitive streak or dissection damage. The remaining 38 embryos were dissected to discard the extraembryonic portion and combined to ensure sufficient number of cells for analysis. The embryos were dissociated into single cells by incubating them in 200μl of TriplE Express for 8 minutes at 37°C with mild mixing every 2 minutes. To stop the TriplE Express, 1ml of ice-cold 10% FBS in PBS was added, and the cells were then filtered through a 40μM Flowmi cell strainer. Following centrifugation at 300g for 4 minutes, the supernatant was removed, and the cells were resuspended in 50μl of PBS containing 0.04% BSA. Cell counts and viability assessments were performed using trypan blue staining on a Countess II instrument (Invitrogen), confirming that over 95% of cells exhibited high sample quality.

The isolation of cell nuclei was carried out following the low-cell input version of the 10X protocol (Protocol Link: https://assets.ctfas-sets.net/an68im79xiti/6t5iwATCRaHB4VWOJm2Vgc/bdfd23cdc1d0a321487c8b231a448103/CG000365_DemonstratedProtocol_NucleiIsola-tion_ATAC_GEX_Sequencing_RevE.pdf). In brief, the 50μl cell suspension was transferred to a 0.2ml PCR tube and centrifuged at 300g for 5 minutes. After removing the supernatant, the cells were resuspended in 50μl of ice-cold nuclear extraction (NE) buffer (containing 10mM Tris pH 7.5, 10mM NaCl, 3mM MgCl2, 1% BSA, 0.1% Tween, 1mM DTT, 1U/ul RNaseIn from Promega, 0.1% NP40, and 0.01% Digitonin) and incubated on ice for 4 minutes. Subsequently,

50μl of wash buffer (similar to NE buffer but without NP40 and digitonin) was added, and the nuclei were pelleted by centrifugation at 500g for 5 minutes at 4°C. Following removal of the supernatant, the nuclei underwent an additional wash with 50μl of diluted nuclei buffer (10x Genomics), were pelleted again, and eventually resuspended in 7ul of diluted nuclei buffer (10x Genomics). A 1μl sample was assessed for quality and nuclei counts using a Countess II instrument, revealing that more than 99% of nuclei stained positively for trypan blue and exhibited the expected morphology. The nuclei were then diluted. A total of 13750 nuclei were taken forward for 10x snATACseq library preparation.

### 10x snATACseq library preparation and sequencing

NGS experiments were performed in the Genomics Unit of the CNIC. Nuclei were counted and their integrity was checked using the Countess III cell counter (Thermofisher). 13750 nuclei were taken for the transposition reaction and loaded into one port of a Chromium Next GEM Chip H (10x Genomics) with a target output of 7,000 nuclei. Single nuclei were encapsulated into emulsion droplets using the Chromium Controller (10x Genomics). Sn-ATAC-seq libraries were prepared using the Chromium Next GEM Single Cell ATAC Kit v1.1 (10x Genomics) following the manufacturer instructions and the library was amplified using a SureCycler 8800 thermal cycler (Agilent Technologies). The average size of the library was then calculated using a High sensitivity DNA chip on a 2100 Bioanalyzer (Agilent Technologies) and the concentration was determined using the Qubit fluorometer (Thermofisher).

Library was loaded at 700 pM onto a P2 flow cell (100 cycles) of the NextSeq 2000 (Illumina) in paired-end configuration (50bp Read1, 8bp Index1, 16pb Index2 and 50bp Read2). FastQ files were obtained using cellranger-atac mkfastq pipeline (10x Genomics).

### snATACseq data processing

Cellranger-atac (v2.1.0) pipeline from 10X Genomics was used to align sequencing data and quantify fragments.

Fragment data was analyzed using Signac (v1.10.0) ^54^ and Seurat (v4.4.0) ^55^ R packages. Cells were filtered using a minimum of 1500 and a maximum of 70,000 reads inside peaks per cell, a minimum percentage of reads in peaks of 15%, a minimum TSS enrichment of 3 and a maximum nucleosome signal (ratio of mono-nucleosome cut fragments to nucleosome-free fragments) of 4.

Doublets were identified using the scDoubletFinder package (v1.12.0) ^56^. Cells were clustered using LSI dimensional reduction and removing the first component, which was highly correlated with sequencing depth.

Cells were annotated with the Mouse Gastrulation Atlas (EmbryoTimecourse2018) ^5^ using the TransferData function from Seurat. Epiblast cells were subsetted and reclustered. Marker peaks were obtained using the FindAllMarkers function and logistic regression. Motif annotations were obtained from JASPAR 2022 database ^57^, and marker peaks were queried for enriched motifs using the FindMotifs function. Motif activities for each cell were calculated using ChromVAR v1.20.2 ^28^.

Topic scores were calculated by finding the overlapping peaks between our dataset and the cell-type specific peaks described in ^58^ for each topic and using those features as input for the AddModuleScore function from Seurat.

### 4D Migration behaviour analysis

#### Data Extraction and Preprocessing

Cell tracking involved extracting data from H2B:miRFP703 embryo live imaging stacks ^40^, focusing on early embryonic progenitors migrating towards the cardiogenic region at the anterior side of the embryo. We obtained 17,170 spatial and temporal coordinates of cells from one embryo after filtering out instances with invalid or missing values. They correspond to tracks of 173 unique cells at the endpoint.

We first generate complete independent tracks for each final cell to prevent assignment problems in cell division scenarios, needed for further processing and analysis. Consequently, each unique cell trajectory identifier encompasses all positions of that cell and all its antecedent cells.

#### Computation of Cell Behaviour Signatures

Cell migration parameters were estimated from cell tracks using CelltrackR v1.1.0 package ^59^ in R, extracting multiple kinetic measurements ^60^. These measurements include track length, displacement (Euclidean distance between start and end points), maximum displacement (from the starting point to any other point within the track), speed, displacement ratio (displacement divided by max displacement), outreach ratio (max displacement/trackLength), straightness (displacement/trackLength), asphericity (similar to straightness but robust to noise by using principal components), overall track angle and dot product measured between first and last segment of tracks, mean turning angle, and fractal dimension (measurement of irregularity). Additionally, we incorporated an additional context-dependent parameter, with measures the distance to the endoderm. We used Matlab R2022b to calculate the distance of each tracked cell to the endoderm, using Matlab wrappers for reading and handling images in KLB format ^61^. To minimize the computational burden of image processing, each time-instant 3D volume was loaded and processed separately, and rescaled to obtain an isotropic image with the existing Z-axis resolution. After segmenting the endoderm by thresholding, a Euclidean distance transform was applied and the distance values corresponding to the positions of the tracked cells in that volume were extracted and stored for subsequent analysis. The distance of a subtrack was defined as the running mean of all the distances.

To better describe the motility behaviour of the cells, and minimize the impact of tracking noise, we decided to calculate these descriptors on each cell across multiple temporal windows of varying sizes. To accomplish this, we constructed all possible subtracks of size w (where w represents the number of timepoints or steps) for each cell, and compute the entire set of measurements described above. Each window w represents a smoothed version of the local behaviour of each cell, ranging from w=1 (5 minutes, instantaneous but possibly noisy measurements), to w=40 (smoothed measurements obtained in cells tracked for 40 frames, 200 minutes). It should be noted that as window size increases, the number of available timepoints per cell decrease, since we cannot create subtracks of length w starting in the last w timepoints.

Collectively, the set of all smoothed versions for the 15 parameters (kinetic and distance to endoderm) conform the behaviour signature of each cell at a specific timepoint.

#### Statistical Analysis of Cell Behaviour Signatures

To test whether there is a difference in behaviour over time between two cell types (CM vs ET) we used linear mixed-effects models. Each temporal response variable was modelled in terms of time, cell type, and their interaction, also incorporating random effects for longitudinal data, and natural cubic splines (with 5 degrees of freedom) to model the potential non-linearities. The interaction terms between cell type and timepoint allow us to check whether the change in the variable over time differs between two groups in each of the 5 time slots into which we divided the temporal response. A w = 70 was used in this first analysis to smooth the potential differentiation delays between cells fated to the same fate. R packages lme4 v1.1.32 ^62^ and lmerTest v3.1.3 ^63^ were used to perform this analysis.

#### Unsupervised Analysis of Behaviours

Behaviour signatures of each cell and timepoint with parameters measured for w=1,5,10,…,40 were first scaled (z-score normalization). N principal components determined by automated elbow point detection on explained variance were used for further clustering with Louvain algorithm, and subject to tSNE dimensionality reduction using Seurat v4.2.1 ^64^ in R.

## SUPPLEMENTARY FIGURES

**Figure S1_1:**
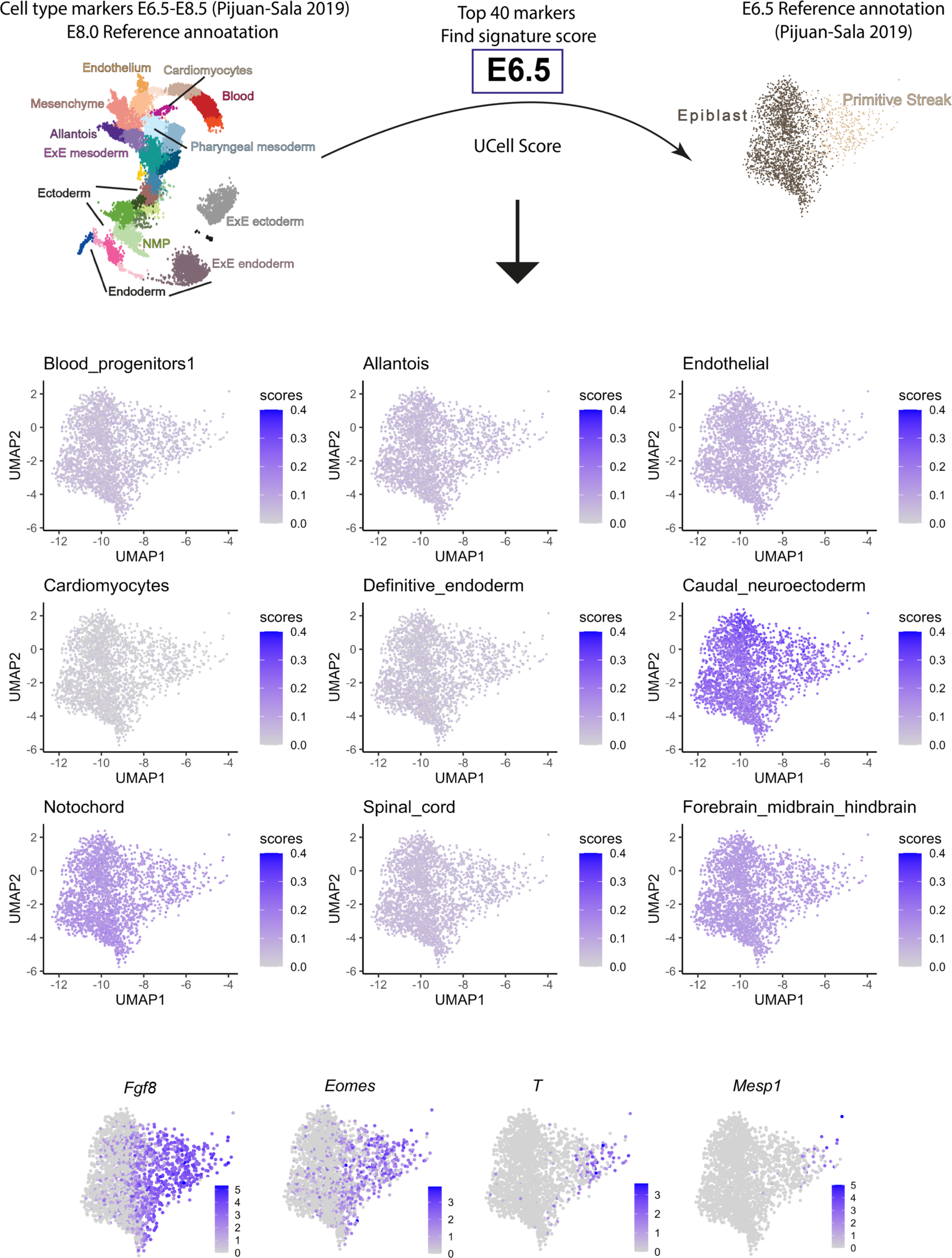
Gene signatures for established cell types at E8.0-E8.5 plotted in E6.5 data. Primitive streak markers gene expression is depicted at the bottom. scRNAseq data from^5^.

**Figure S1_2:**
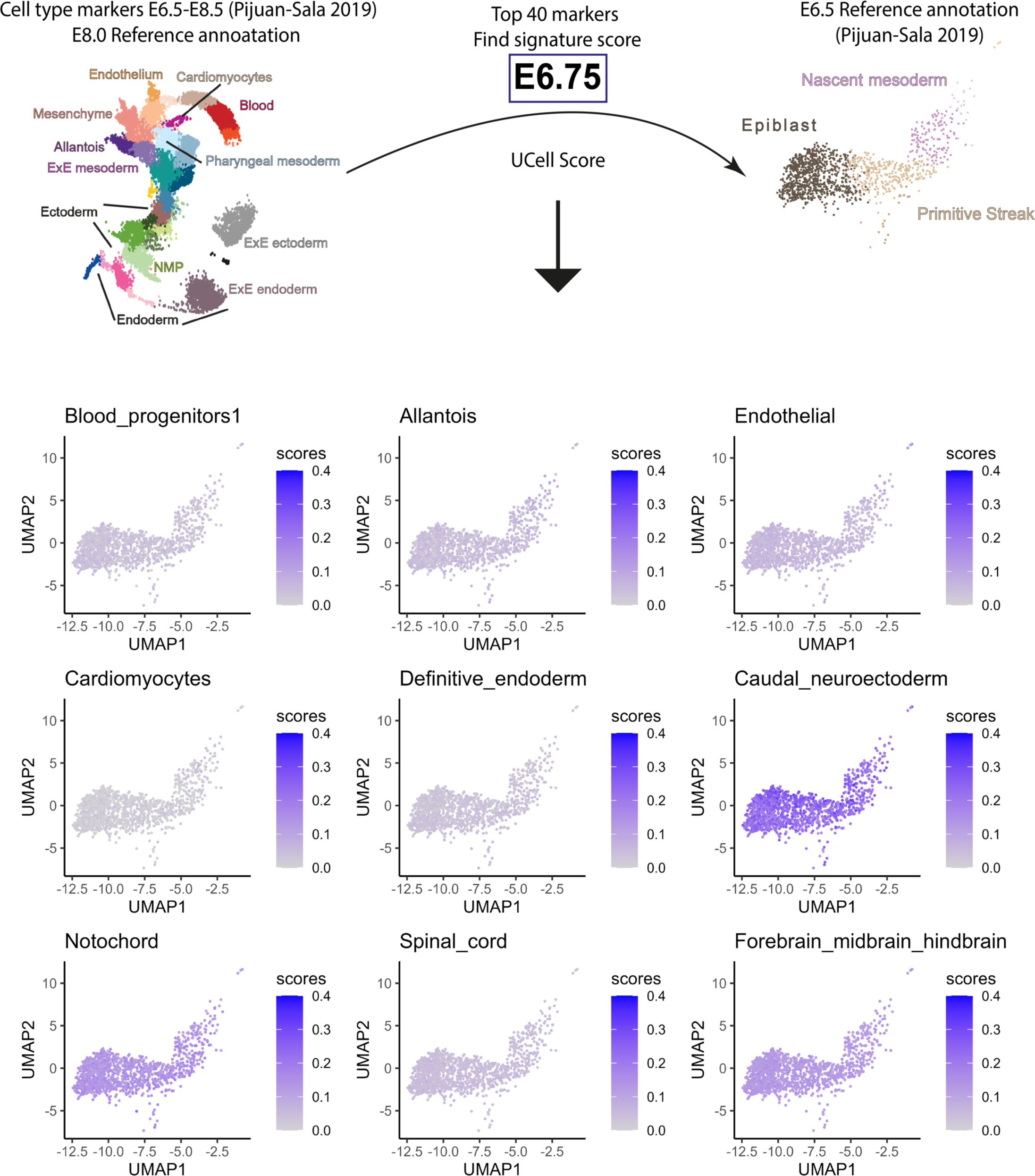
Gene signatures for established cell types at E8.0-E8.5 plotted in E6.75 data. scRNAseq data from^5^.

**Figure S1_3:**
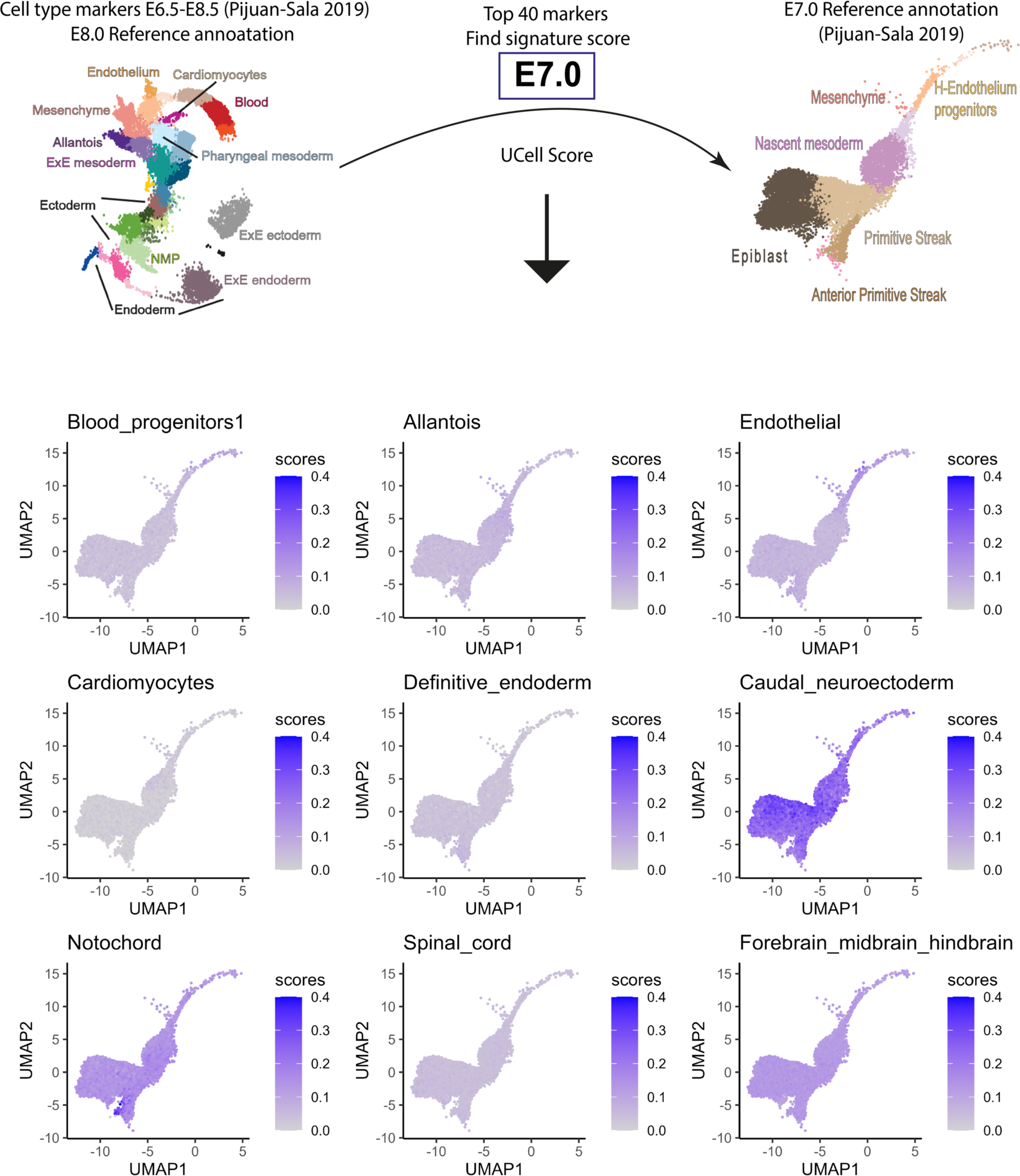
Gene signatures for established cell types at E8.0-E8.5 plotted in E7.0 data. scRNAseq data from^5^.

**Figure S1_4:**
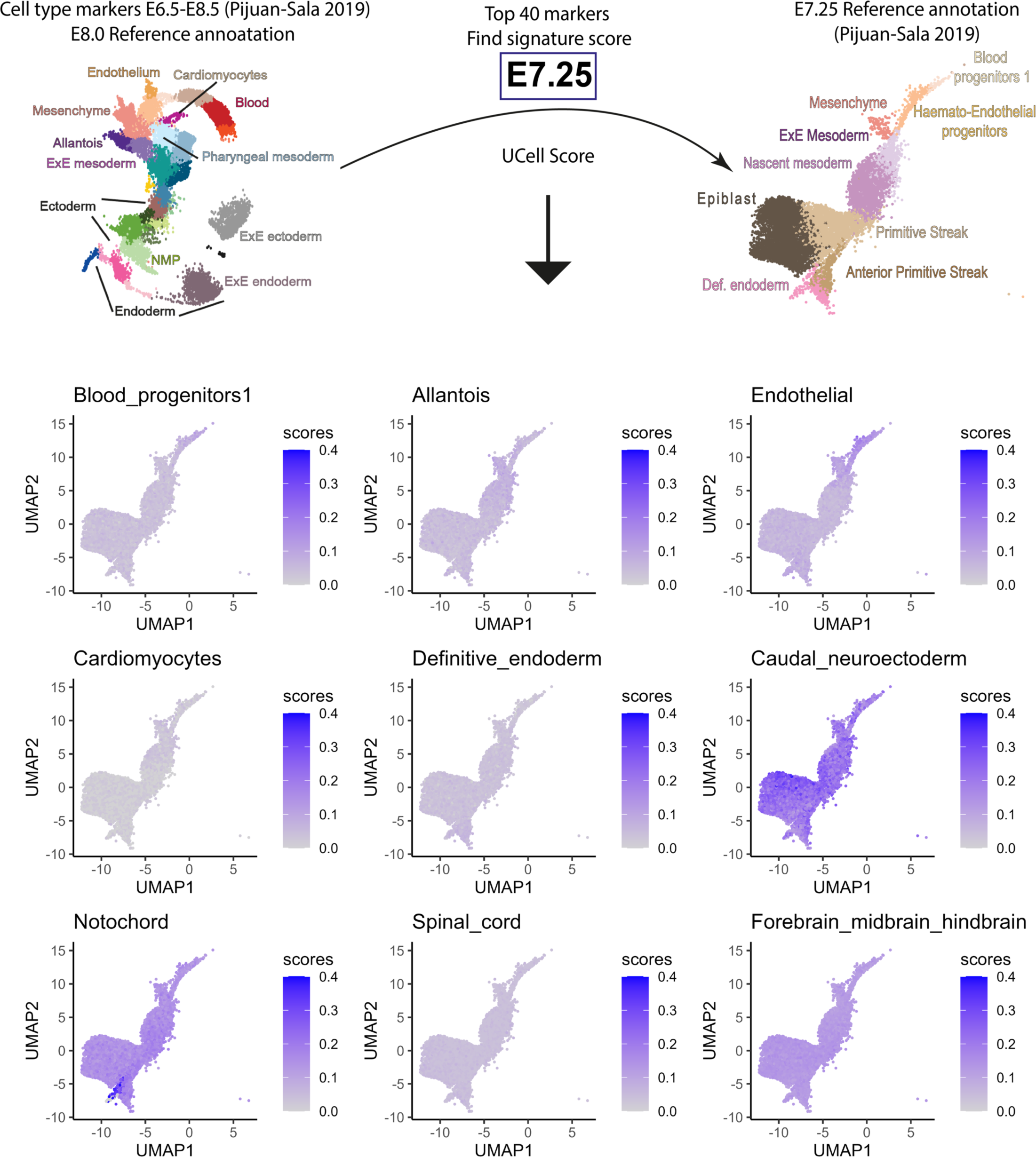
Gene signatures for established cell types at E8.0-E8.5 plotted in E7.25 data. scRNAseq data from^5^.

**Figure S1_5:**
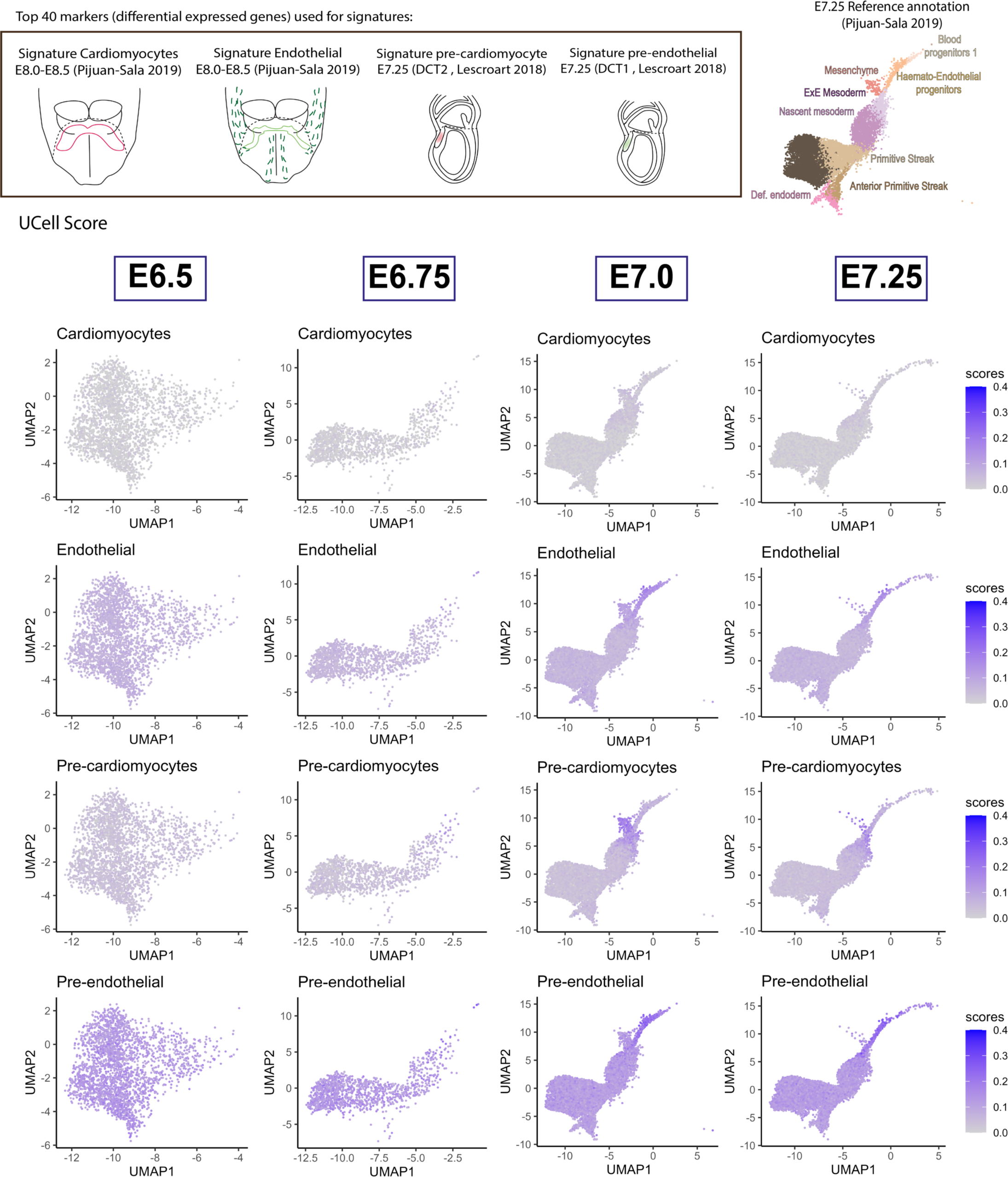
Emergence of Cardiomyocyte and endothelial expression signatures from E6.5 to E7.25. Top: signature scores based on “Car- diomyocyte” and “Endothelium” markers from^5^. Bottom signature scores based on “DCT2- Pre-Cardiomyocyte” and “DCT1-Pre-Endothelium” markers from^24^. Both plotted on data from^5^ .

**Figure S2:**
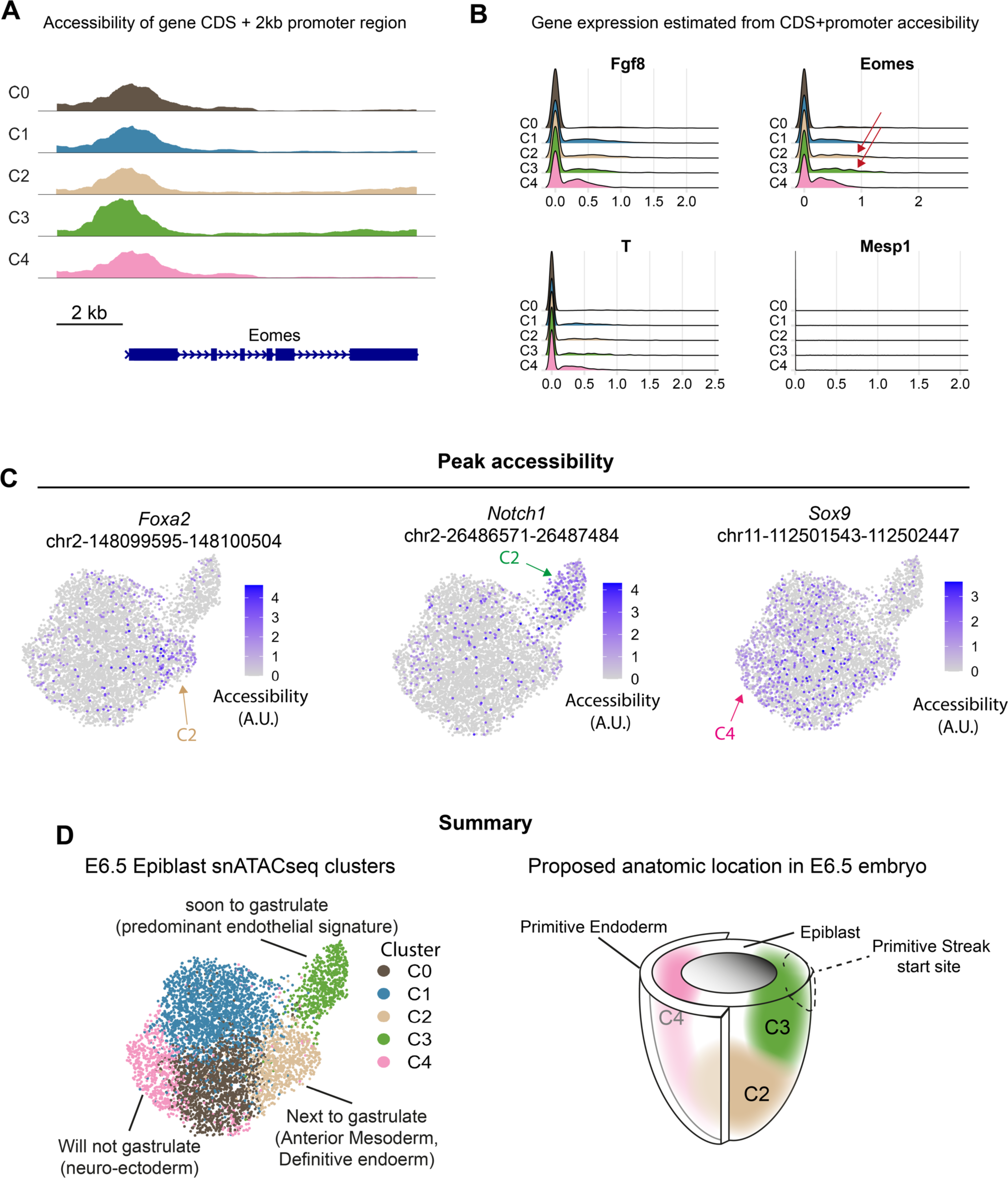
Predicted expression of gastrulation regulators. (A) Coverage plot of ATAC seq reads for *Eomes* coding sequence and promoter. Each cluster is represented as a pseudo-bulk. (B) Inferred RNA expression of genes expressed during the onset of gastrulation. Red arrows point to *Eomes* expressing cells. (C) Accessibility of peaks associated to genes that are typically expressed in definitive endoderm (*Foxa2*), endothelial (*Notch1*), and ectodermal progenitors (*Sox9*), respectively. (D) Epigenetic priming for snATACseq clusters in the E6.5 epiblast and their proposed location in the embryo.

**Figure S3:**
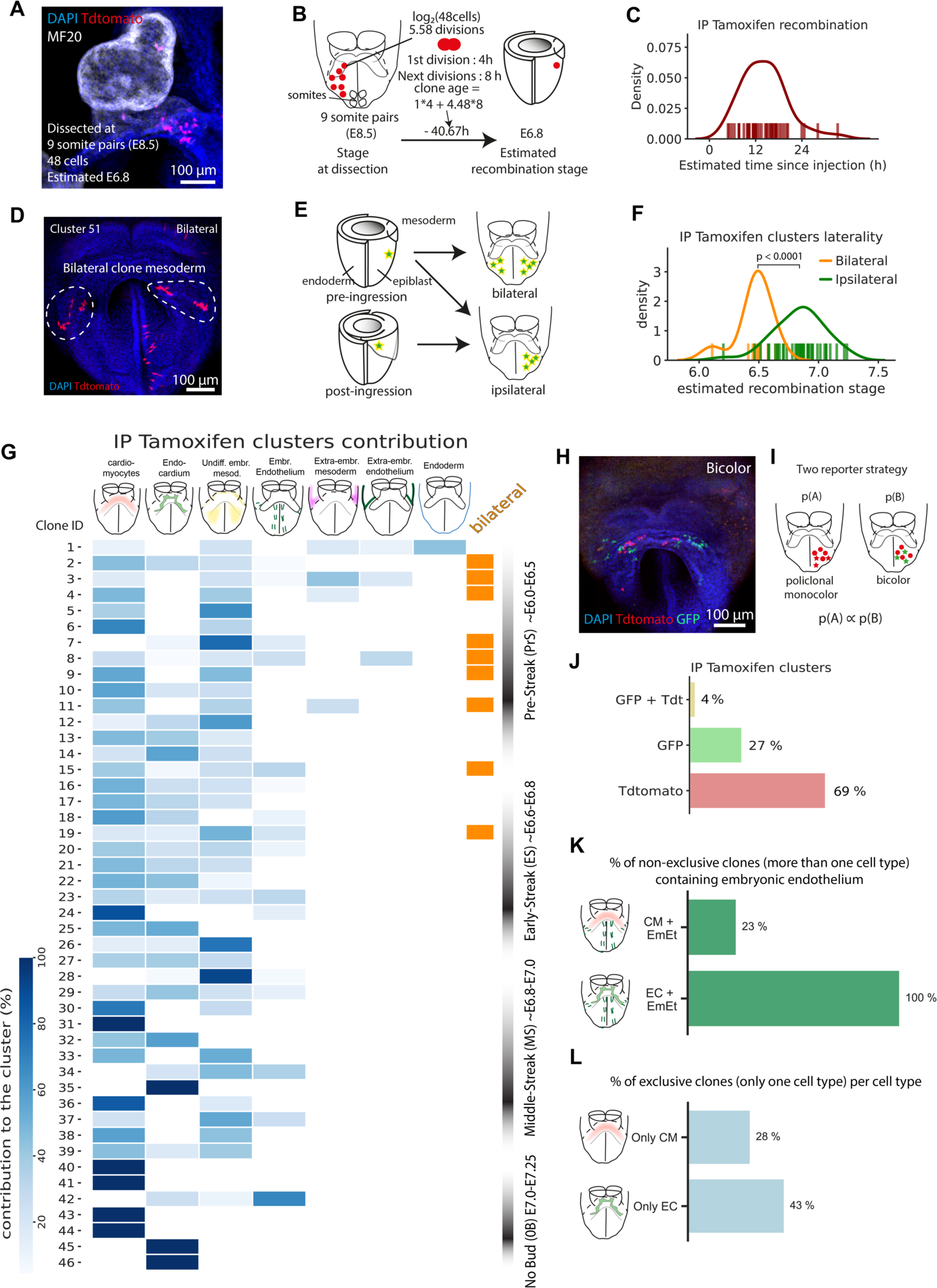
**Estimation of the stage of clone recombination by cell count and detailed contribution of 4–OH Tamoxifen induced clones. (**A) Intensity projection of a confocal image showing a clone and its estimated recombination stage. (B) Rationale of the estimation of recombination time from embryo stage and cell number. (C) Kernel density estimate revealing the distribution of the estimated delay in recombination since tamoxifen injection, and rug plots at the bottom showing data for individual clones (n = 44 embryos). (D) Intensity projection of a confocal image showing a bilateral clone in the mesoderm. (E) Recombination events that occur in cells before ingression to the mesoderm can give rise to either bilateral or ipsilateral clones, while post–ingression events can only give ipsilateral clones. This serves as an internal reference for the time estimation method. (F) Kernel density estimate revealing the distribution of estimated recombination stages of the entire cluster collection, and rug plots at the bottom showing data for individual clusters (n = 44 embryos, 46 clones of which 9 are bilateral and 37 unilateral). A Mann-Whitney U rank test on two independent samples was performed to compare both distributions. (G) Heat map showing the contribution of each clone. Each square contains a cell count for every clone (X–axis) and location (Y–axis), and is color graded for its contribution weight to the clone. Bilateral clones are marked with an orange square. Estimated stage at recombination is shown on the right. (H) Example of a bicolor cluster (n = 2 embryos). (I) The rationale of the two–reporter strategy. (J) Percentage of bicolor, GFP and Tdtomato clusters in the clonal analysis collection (n = 44 embryos). (K) Percentage of clones cardiomyocyte+other mesoderm clones containing other endothelial cells (not endocardial) versus percentage of endocardium + other meso- derm clones containing other endothelial cells. (L) Percentage of exclusive cardiomyocyte or endocardial cells (not containing any other cell type).

**Figure S5:**
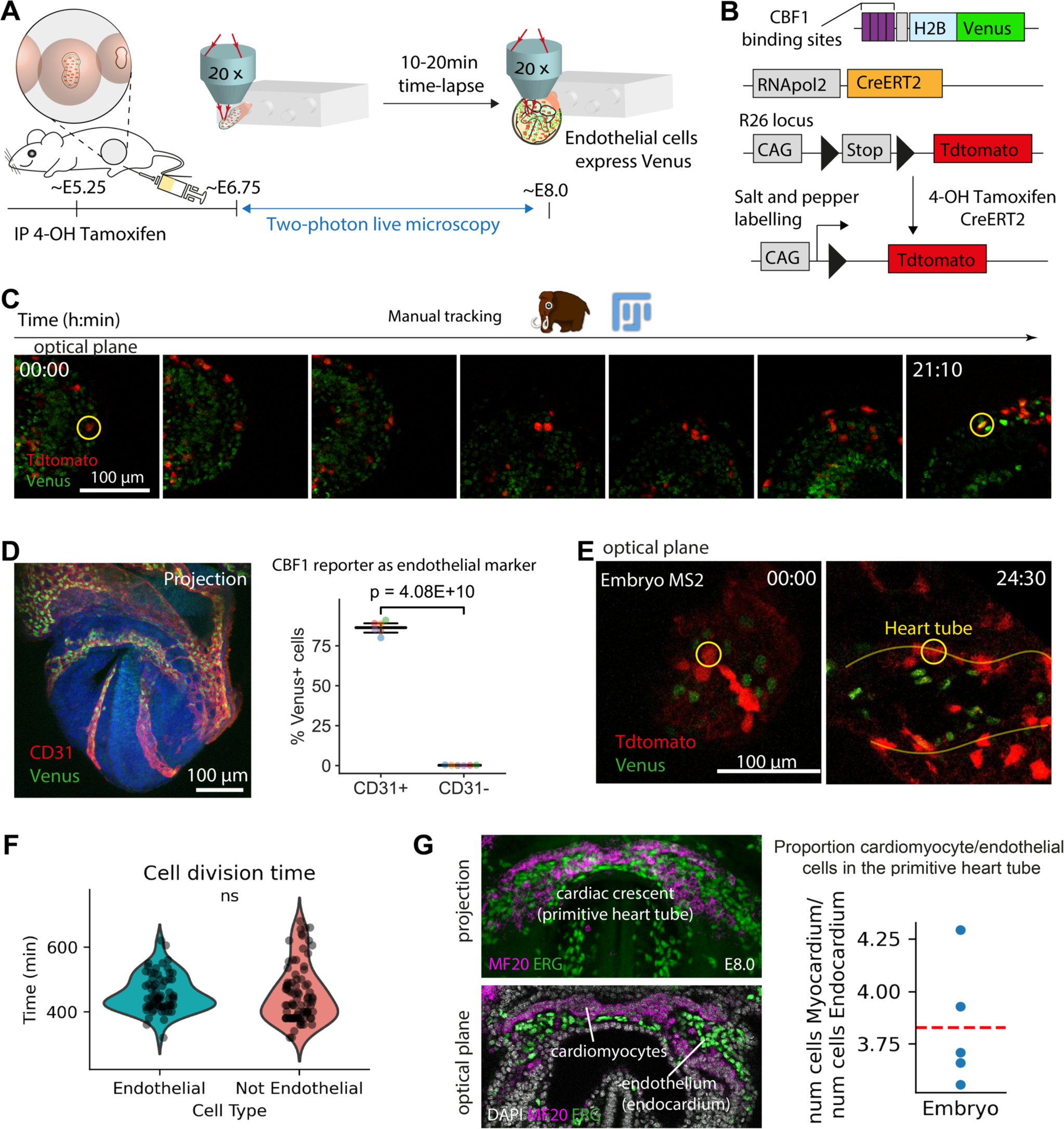
Two–photon time–lapse microscopy for cardiac progenitor cell tracking and lineage reconstruction. (A) Diagram of the experi- mental setup in the two–photon microscope. (B) A NOTCH activity reporter^44^. allows endothelial nuclei identification. Random induction of the Tdtomato reporter allows cell tracking. (C) Selected time points for the tracking of an endocardial progenitor from the nascent mesoderm to the primitive heart tube. (D) Venus (NOTCH) positive cells are identified as endothelial cells by CD31 immunostaining (n = 6 embryos, t–test for two related samples). (E) Initial and final time points of a cardiomyocite progenitor track in a RERT;tdtomato H2B:Venus embryo. (F) Time between tracked cell divisions in endothelial and not endothelial cell progenitors (n = 151 divisions, 3 embryos, t-test two related samples). (G). Whole mount immunostaining of E8.0 embryos to count the number of MF20+ (cardiomyocytes) and ERG+ (endothelial cells) at the primitive heart tube, which ratios are shown for every embryo in the right panel. A dashed red line marks the average at 3.8.

**Figure S6:**
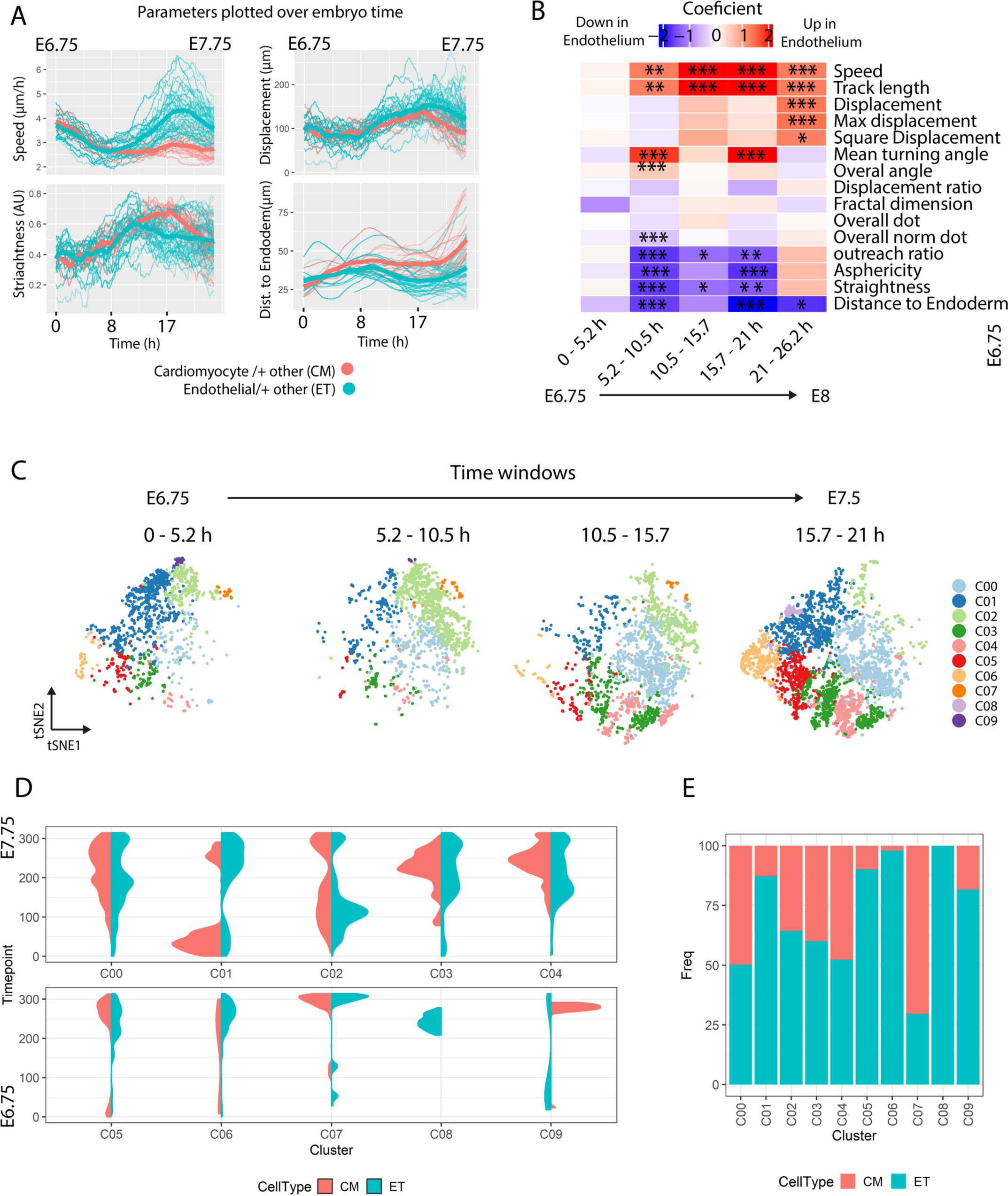
**Temporal diversification of migration behaviour in early cardiac progenitors. (**A) Univariate analysis of longitudinal data, investigat- ing differences in average responses over time between cell types endothelial and not endothelial, using a linear mixed-effects model with coefficients providing insights into the trends and variation in 5 time windows. (B) Temporal evolution of migration parameters in endothelial and non-endothelial cardiac progenitors, single tracks in thin line, average in thick line. Data in both panels A and B represent a 350 min moving window. (C) UMAP plots representing clusters segregated by time windows. (D) Violin plots for the normalized distribution of timepoints for each cluster and endpoint cell fate. (E) Proportions of endpoint cell fate in each cluster.

